# The Antimicrobial Gut Resistome of the Wayampi reveals a shared *background* of antibiotic and metal resistance genes with industrialized populations, underscoring the “robust-yet-fragile” architecture of human gut microbiomes

**DOI:** 10.1101/2025.06.30.662145

**Authors:** Miguel D. Fernández-de-Bobadilla, Ana Elena Pérez-Cobas, Antoine Andremont, José Luís Martínez, Fernando Baquero, Val F. Lanza, Teresa M. Coque

**Author notes:** Corresponding author: Teresa M. Coque. Servicio de Microbiología, Hospital Universitario Ramón y Cajal, Carretera de Colmenar, km. 9.1, Madrid 28034, Spain. Co-corresponding author: Val F. Lanza. Unidad de Bioinformática, Hospital Universitario Ramón y Cajal (IRYCIS), Carretera de Colmenar, km. 9.1, Madrid 28034, Spain.

## Abstract

**Background:** Metagenomics enables detailed profiling of antimicrobial resistance genes, but most studies focus solely on antibiotic resistance genes (ARGs), use low-sensitivity methods, and rarely investigate populations with minimal anthropogenic pollution. We analyzed fecal resistomes of 95 Wayampi individuals, an Indigenous community in remote French Guiana, using a targeted metagenomic capture platform covering 8,667 resistance genes to antibiotics (ARGs), metals (MRGs), and biocides (BRGs) (PMID: 29335005). Resistome profiles were compared with Europeans to assess population-level differences.

**Results:** ARG richness was similar between groups (259 in Wayampi *vs*. 264 in Europeans, 159 shared), but MRGs + BRGs (BAC) gene richness was signifcantly higher in Wayampi (3,670 vs. 2,039). Most genes appeared in a minority of individuals (mean 5% for ARGs, 2% for BAC), but several ARGs for tetracyclines [*tet(32), tet(40), tet(O), tet(Q), tet(W), tet(X*), *tetAB*(P)], aminoglycosides (*ant6*′*-I*, *aph3-III*), macrolides (*ermB*, *ermF*, *mefA*), and sulfonamides (*sul2*) were present in all individuals. Tetracycline resistance genes predominated overall, while beta-lactam resistance genes were more common in Wayampi, and genes conferring resistance to aminoglycosides, amphenicols, and folate inhibitors were more frequent in Europeans. Among MRGs, copper and arsenic resistance genes prevailed, followed by those for zinc, iron, cobalt, and, nickel. Up to 76% of Wayampiis carried acquired MRGs for copper (*pcoABCDRS* and *tcrB*), silver (*silACFPRS*), arsenic (*ars*), and mercury (*mer*) detoxification. Shannon diversity indices were similar for ARGs, MRGs, and BRGs, but composition and evenness differed significantly. UMAP and ADONIS analyses distinguised cohorts by ARG profiles (*p* < 0.001), but not MRGs or BRGs. Correlation analysis revealed conserved gene-sharing networks and introgression of acquired ARGs and MRGs in both gut microbiomes.

**Conclusions:** The diverse and balanced Wayampi resistome reflect a less perturbed microbiome compared to industrialized populations, and reveals a *background* of core and shell acquired ARGs and MRGs, consistent with the “robust-yet-fragile” architecture of scale-free networks. The patchy yet resilient distribution of resistance genes reflects varying levels of conserved gene sharing highways probably shaped by long-term microbial-human evolution, and support a broader view of acquired antimicrobial resistance.

## BACKGROUND

Antimicrobial resistance (AMR) is a major global health challenge that must be approached through the lens of *disease ecolog*y [1]. Understanding the interactions between microbes, hosts, and their environments and how these interactions shape the spread and evolution of AMR, is essential for identifying and managing hotspots where resistant populations and genes emerge, concentrate and evolve [2]. Aditionally, it can help to inform efficient strategies to mitigate the selection, mobilization, and persistence of resistance determinants [3–5].

The evolution of antibiotic resistance can be broadly divided in two ages [6]. Intrinsic antibiotic resistance is a natural and ancient phenomenon that mirrors the microbial phylogeny [7] while acquired antibiotic resistance emerged in more recent times driven by the therapeutic use of antibiotics. This has led to the rapid spread of a relatively small number of acquired antibiotic resistance genes (ARGs) partially decoupled from the overall evolutionary trajectories of bacterial species. All bacteria harbour intrinsic resistance genes, many of which have a functional role beyond mere antimicrobial resistance [8,9]. In contrast, the diversity and abundance of acquired ARGs seem to be largely shaped by anthropogenic selective pressures and the exchange of transferable ARGs within and across microbiomes [1,4,10–13].

Several works have examined the antibiotic resistomes of humans, animals, and environmental samples [12,14,15]. However, interpreting and comparing results across these studies are often challenging due to technical limitations inherent in metagenomic methods and biases in sampling design. Firstly, the total number of ARGs identified in most metagenomic studies is often low, largely due to the limited sensitivity of currently used omic approaches. Gene capture platforms significantly enhance the recovery of ARGs but they have been used for resistome analysis in a few studies [16,17]. Secondly, characterizing *background* and *baseline* resistomes is essential for contextualizing the presence of ARGs in specific ecosystems [3,18]. At present, there is an overrepresentation of cross-sectional studies centered on humans in industrialized countries, limiting our ability to differentiate between “*background*” ARGs (those naturally present in an ecosystem) and “contamination” (referring to ARGs introduced and selected by human activity) [18]. Finally, resistance to non-antibiotic antimicrobials, particularly metals and biocides, which are extensively used across various One Health sectors, remains underexplored. This oversight is surprising given the historical and growing significance of metals in therapeutics contexts [19,20] and their relationship with the (co)selection and persistence of antibiotic resistance [21].

In this study, we characterized the antimicrobial resistomes, which include gene pools associated with resistance to antibiotics (ARGs), heavy metals (MRGs), and biocides (BRGs), of the Wayampi, an Indigenous population from a remote geographical community of French Guiana. This community, previously shown to harbour the most diverse human gut microbiome [22], has had minimal exposure to antibiotics or conventional healthcare systems and maintains a non-industrialized lifestyle [23,24]. Despite their geographic and cultural isolation, the Wayampi are chronically exposed to mercury pollution from artisanal gold mining, a widespread activity in the Amazon that affects human health [25–27] and might influence their MRGs profile [30]. These characteristics make the Wayampi an unique model for studying the human antimicrobial resistome. While previous studies on this Amerindian community have provided valuable insights [22–24,28–30], a comprehensive catalog of ARGs remains elusive due to the limited resolution of conventional metagenomic and culturing methods, and their MRGs pool remains largely underexplored. To address this gap, we used ResCap, a highly sensitive and specific targeted metagenomic approach capable of detecting ARGs, MRGs, and BRGs [17]. We integrated our results with existing data, developed a custom bioinformatic pipeline for large scale ResCap-analysis, and examined the influence of demographic variables on resistome composition.

## MATERIALS and METHODS

### Study subjects and collection of samples and data

The study involved 95 Amerindians from the remote Wayampi community in the southernmost region of French Guiana [31]. Their living conditions were described elsewhere. Briefly, the residents inhabit large huts, with an average of 13.9 inhabitants per hut, ranging from 4 to 38 individuals, distributed across three villages located near gold mining areas. The community lacks access to latrines or any sanitary facilities. Their diet primarily consists of local food sourced from traditional crops, fishing, or hunting, with minimal farming activities; only a few free-running chickens are kept. Access to antibiotics is restricted to a health post in one of the villages, where medications are provided free of charge upon prescription by resident nurses. For serious medical needs, villagers can seek treatment at hospitals in Cayenne, the capital of French Guiana. The community has limited interaction with outsiders and is approximately 100 km south of the nearest village. Access to the village is highly regulated by regional authorities. Estimates indicate an average antibacterial use of 0.64 treatments per person per year, and the hospitalization rate is 6.1% per year [23,24].

The Wayampi community exhibits elevated levels of mercury in their hair, with some individuals exceeding the concentration levels recommended by the World Health Organization [25,32]. Demographic information, including age, sex, lifestyle details (diet, daily activities, travel frequency), and the size and location of households, along with medical records, were collected in the year prior to sampling at the health post and through interviews with local informants. Samples from adults in France (n=29), who had no exposure to antibiotics in hospitals, were included as a control group [33]. The collection and processing of samples were conducted as described in previous studies [23,33].

### Targeted metagenomics experimental and bioinformatic workflow

The characterization of resistomes (ARGs, MRGs, BRGs) was conducted using ResCap-targeted sequence panel, which is based on the custom SeqCap EZ library format (NimbleGen, Madison, USA) [17]. ResCap has a target space of 88.13 Mb and contains probes for 78,600 non-redundant genes. This includes 47,806 putative ARGs and 7,963 experimentally characterized genes, alongside 30,794 putative resistance genes and 704 canonical MRGs and BRGs that have been deposited in the CARD, ResFinder, and BacMet databases [34–36]. DNA was extracted from 250mg of fecal samples using the Powerfecal Qiagen (QIAGEN, UK), following the manufacturer’s instructions. The quality and concentration of the extracted DNA were measured using a NanoDrop 2000 Spectrophotometer (Thermo Scientific, Waltham, MA, USA) and a Qubit® 2.0 Fluorometer (Life Technologies, Waltham, Massachusetts, USA). The ResCap workflow consists of (i) whole-metagenome shotgun library construction, (ii) hybridization and capture, and (iii) captured DNA sequencing. All steps were performed in our institution’s facilities (https://www.irycis.org/es/servicios/11/genomica-traslacional-y-bioinformatica) according to the standard protocols provided by Nimblegene for Illumina platforms [17].

The raw captured on-target sequences were analyzed using the sequence aligner bowtie2 [37]. Further accurate gene identification consisted of using KMA, a robust mapping method designed to map raw reads directly against redundant databases and distinguish between gene alleles in a sample [38]. As a reference database, we built a custom-made library based on the latest version of Resfinder [39] and BacMet [36]. We also developed a pipeline to transform the gene data from BacMet initially presented in the protein version to nucleotide sequences. We removed genes with a p-value higher than 0.05 and an identity lower than 90% for ARGs, MRGs and BRGs, in our custom database. Due to the extensive BacMet database size, the identity threshold for the genes included here was 90%. To normalize the results, we performed a rarefaction analysis using the “rrarefy” function from the package vegan implemented in R.

### Ecological analyses

We compared the number and diversity of genes between Wayampi and European populations using the Mann-Whitney test and further visualized the gene distribution among groups using the “ggstatsplot” package. A pseudo-rarefaction test was conducted to validate the sampling size, as outlined by Colwell et al. [40]. Next, we categorized genes according to their frequency distribution into three groups, namely, “core genes” (present in ≥ 90% of individuals within a given cohort), “high prevalence genes” or “shell genes” (present in 15-80% of individuals), and “low prevalence genes” (present in less than 15%). This classification is based on the earlier work of Wolf and Koonin regarding pangenomes [41]. To assess the significance of the differences in the proportion of each gene across cohorts, we applied the chi-square test. Additionally, we employed a general linear model (GLM) to analyze the differential abundance of genes, focusing on those with a p-value lower than 0.05 between cohorts. To visualize the results, we created volcano plots and heatmaps using the R package pheatmap [42].

Alpha diversity was quantified using the Shannon index, Chao 1 richness estimator, and Simpson index, all implemented in the R package “phyloseq”. The normality of the resulting values was assessed through the Shapiro test from the base Rstats package. To compare the distribution of diversity metrics between groups, we used the Wilcoxon test for non-normally distributed data and the Student’s t-test for normally distributed data. Various dimensionality reduction algorithms, including Principal Component Analysis (PCA, from the Rstats package), Uniform Manifold Approximation and Projection (UMAP), and t-distributed Stochastic Neighbor Embedding (t-SNE, from the Rtsne package), were employed to identify compositional patterns and clustering among samples. To assess significant differences in resistome composition between groups, we performed a permutational multivariate analysis of variance (PERMANOVA) using the “adonis2” function from the R vegan package [43].

The impact of demographic variables on the gene composition of fecal samples of each cohort was analyzed using the adonis2 function. This method identifies genes with statistically significant differences in abundance across various sample groups. All analyses were conducted using R software (version 4.3; R Foundation for Statistical Computing, Vienna, Austria).

### Binning of resistome datasets

Correlations within and between the ARGs, MRGs, and BRGs categories were assessed using the rcorr function (Spearman distribution), from the Hmisc R package [44]. Significant correlations were defined by an absolute Spearman’s ρ > 0.5 and a Benjamini– Hochberg adjusted p-value < 0.001. A correlation-based network was then constructed using the Gephi software and the ForceAtlas2 algorithm [45].

## RESULTS

### The ResCap pipeline

We analyzed the resistomes of a Wayampi population using ResCap, a targeted metagenomic platform developed by our group [17]. For this study, we improved the analysis pipeline to efficiently process large datasets. The automated workflow converts Illumina raw data into a tabulated outputs, significantly accelerating bioinformatic analysis, and improving speed, computational scalability, and precision. It support flexible normalization, including rarefaction and mean-based normalization, and incorporates automated outlier detection and exclusion to ensure data quality. The pipeline is freely available on GitHub (https://github.com/migueldiezfdz/ResCap_analysis). A full summary of statistical analysis and results is summarized in **Table S1**.

### Distribution of genes encoding antimicrobial resistance (ARGs, MRGs, BRGs)

To ensure the suitability of the dataset, we first performed rarefaction analyses by gene category **(Figure S1)**. The rarefaction curves plateaued for the Amerindian sample, indicating that the number of samples in this group was sufficient to accurately estimate resistome diversity. Next, we examined the *distribution* of each gene category. **Figure 1A** illustrates that both Wayampi and European samples had a similar number of ARGs (259 vs 264) but differed in the number of MRGs and BRGs (here, considered together as “BAC” because some genes are classified as both MRGs and BRGs in databases; 2039 in Europeans vs. 3670 in Wayampi). Genes were unevenly distributed, with many shared between both populations (**Figures 1B)** and some found in all individuals, representing core genes (**Figure 1C**). Specifically, we identified 103 ARGs and 2,390 BAC only in Wayampi, and 109 ARGs and 759 BACs only in Europeans (representing 40% of of ARGs in each group and 66% and 37%, respectively. **Figure 1D** shows a detailed analysis of the evenness (abundance of each gene) within each gene category in Wayampi and Europeans. We found that most genes were present in a low number of individuals (mean of 5-6% and 2,1-3,4% of the cohorts for ARGs and BAC, respectively), with specific ARGs present in all individuals of each group. (p-values < 0.05 for the three categories, Mann-Whitney test). **Figure 1E** shows that Europeans had a higher average number of ARGs and BACs. However, the median number of genes per person was not significantly different between populations. (p-value > 0.05; Mann-Whitney test). We also categorized the genes based on their frequency distribution into “core genes” (present in ≥ 90% of individuals within a given cohort), “high prevalence genes” or “shell genes” (89-20%), and “low prevalence genes” (<20%). **Figures 1F, 1G,** and **1H** represent gene frequency distributions and show the typical U-shaped pattern described in pangenome models by Koonin and Wolf, in which core and low-prevalence genes are the most abundant [41].

**Figure 1.**
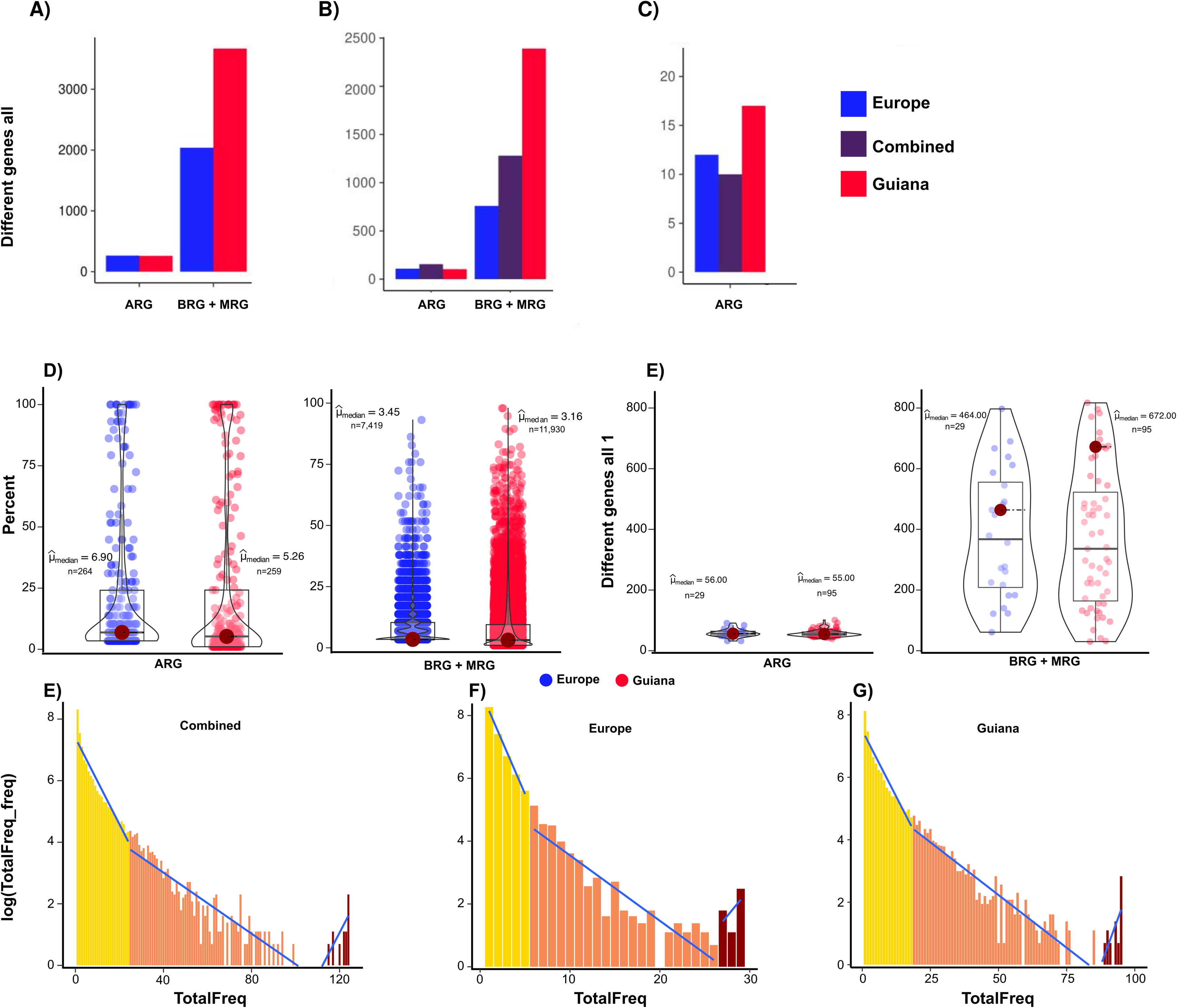
Distribution of genes for Amerindian and European populations. A) Number of unique genes in each gene category for each population. B) Total number of genes that appear at least once in each population and across both cohorts, grouped by gene category. C) Number of core resistance genes classified by gene category and cohort. D) Frequency distribution of each gene within each population. E) Median number of resistance genes per individual in each cohort. F) Histogram of gene frequencies in both cohorts. G) Histogram of gene frequencies in the European cohort. H) Histogram of gene frequencies in the Wayampi cohort. ARG=antibiotic resistance gene; BAC=MRGs (metal-resistance genes) + BRG (biocide-resistance genes).

### Richness and abundance of gene categories (ARGs, MRGs, BRGs)

We conducted a further analysis of the richness (the number of distinct genes) and the relative abundance of different subcategories within the ARGs, MRGS, and BRGs pools of each population (**Figure 2**). In the ARGs category, we found that genes associated with tetracycline resistance were the most prevalent in both cohorts (p-value < 0.05; Mann-Whitney test). However, genes conferring resistance to beta-lactam resistance were prominent in Amerindians, while genes coding for resistance to aminoglycosides, amphenicols, and folate inhibitors were more frequent in Europeans (p-values < 0.05; Mann-Whitney test) (**Figure 2A**). Regarding MRGs, genes associated with resistance to copper and arsenic were the most common, followed by those for zinc, iron, cobalt, and nickel in both cohorts. Differences in the prevalence of genes linked to copper, silver and mercury were observed (**Figure 2B and 2D**). Although the distribution of BRGs was also similar in both cohorts, genes with activity against quaternary ammonium salts were more abundant in Amerindians, whereas genes related to xanthenes were more prevalent in Europeans (p-values < 0.05; Mann-Whitney test) (**Figure 2C**). A detailed analysis of the pools of ARGs, MRGs, and BRGs is given below.

**Figure 2.**
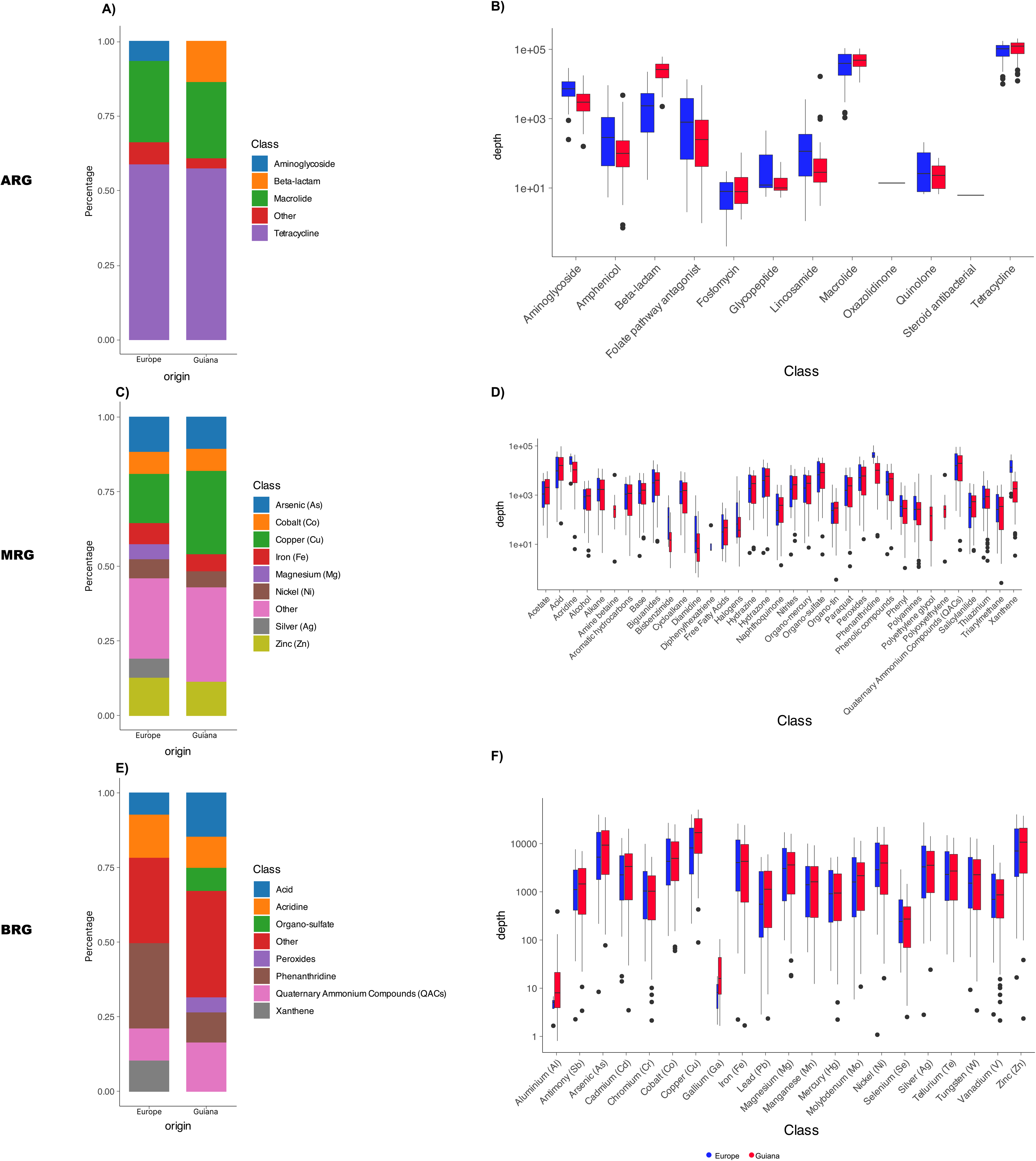
Gene diversity and gene depth across resistance categories. Panels A), C), and E) Proportion of ARGs, BRGs, and MRGs across the population, displayed on a logarithmic scale to highlight relative gene abundance and diversity. Panels B), D), and F) Sequencing (read) depth for each resistance gene category, also shown on a logarithmic scale to illustrate differences in gene representation and coverage. ARG=antibiotic resistance gene; BAC=MRGs (metal-resistance genes) + BRG (biocide-resistance genes).

### Diversity and distribution of ARGs (the antibiotic resistome)

The distribution and diversity of ARGs across the most relevant antibiotic families and cohorts is illustrated in **Figure 3** and **Figure S2**, and supported by **Table S2**. The pool of ARGs in both Wayampi and European populations is similar in size (259 and 264, respectively), with more than half of these genes being present in both groups (156 ARGs). These ARGs provide resistance to various families of antibiotics, including tetracyclines (n=24 genes, 43 allele variants), macrolides (n=16 genes and 35 allele variants), beta-lactams (29 gene categories and 116 allele variants), aminoglycosides (31 genes and 53 allele variants), phenicols (12 genes), quinolones (3 genes and 12 allele variants), glycopeptides (n=4 genes), and fosfomycin (n=5 genes). Most genes were detected in a minority of individuals (mean 5% for ARGs, 2% for BAC). Some genes, referred to as “core genes”, were present in all individuals, while others were found sporadically. The distribution of “canonical (acquired) ARGs” varied significantly among individuals.

**Figure 3.**
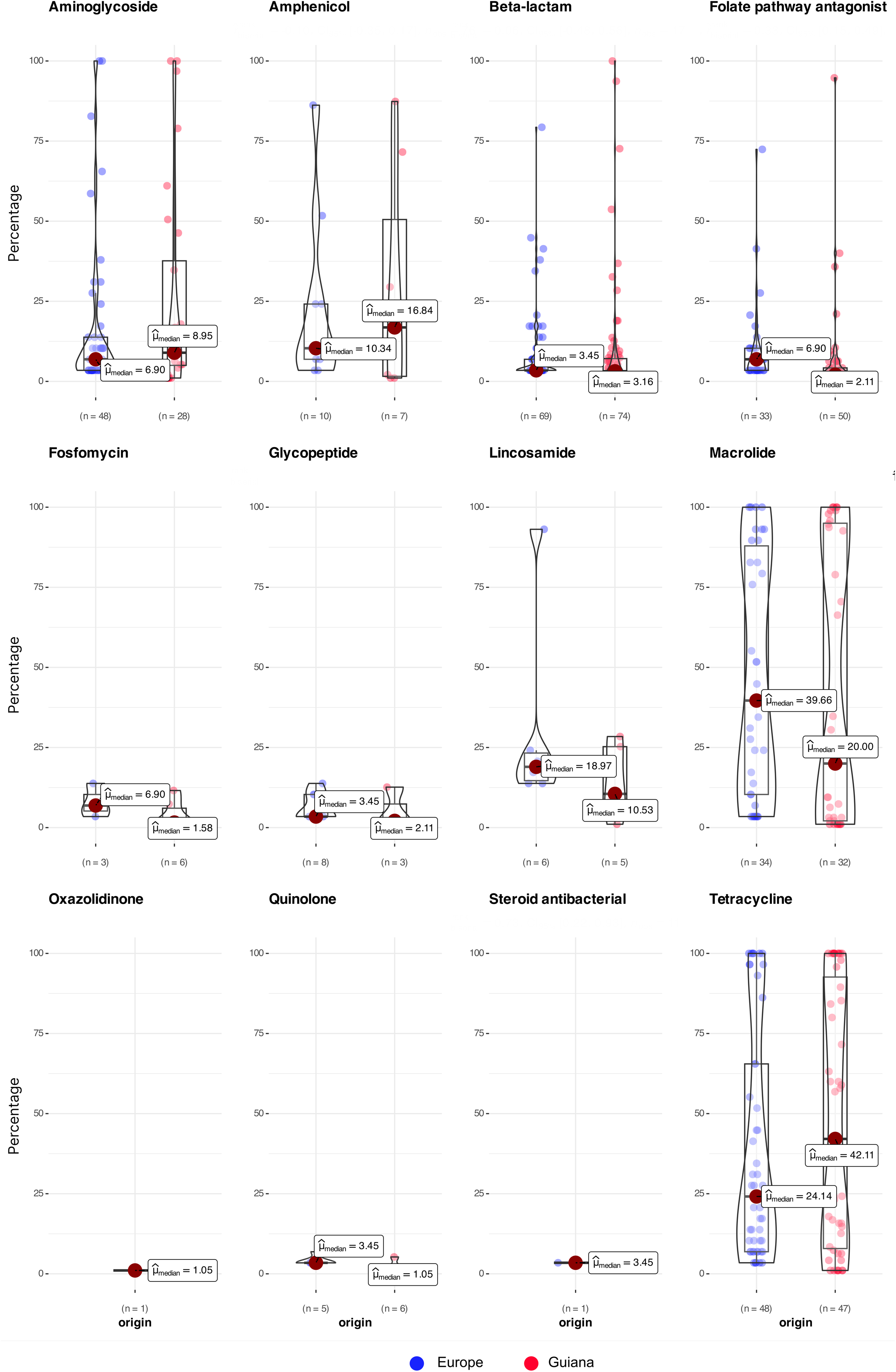
Distribution of antibiotic resistance genes (ARGs) by antibiotic class and population. Each dot represents a specific gene, grouped by antibiotic family. The x-axis indicates the different cohorts, while the y-axis shows the prevalence of each gene, expressed as the percentage of individuals in which the gene is detected.

The “core genes” confer resistance to various antibiotic families, including tetracycline [*tet(32), tet(40), tet(O), tet(Q), tet(W), tet(X*), *tetAB*(P)], aminoglycosides (*ant6’-I, aph3-III*), macrolides [*erm(B), erm(F),* and *mef(A)*] or sulfonamides (*sul2*). Notably, most of these genes have been associated with mobile genetic elements (MGEs) [46,47]. For example, the conjugative transposon (CTn)DOT comprises *tet(Q)* and *ermF* and is found in phylogenetic distant taxonomical groups of the Bacteroidia class such as the Bacteroidales order (Bacteroides, Prevotella genera), or Flavobacteriales (Capnocytophaga genus). The gene *erm(B)* is located on a plethora of composite and conjugative transposons added Clostridiales to the previously mentioned taxa. Additionally, the *mef(A), msr(D)*, *aph3-III,* and *ant6’-I* are located on a variety of transposable elements within Bacillales and Lactobacillales (mostly streptococci, enterococci, and staphylococci). There are also other less common ARGs that provide clinical resistance to different antibiotic families, typically located on MGEs such as class 1 integrons (*dfrA1, sul1, aadA1*) and transposable units of enterobacterales [(*aph(3’’)-Ib aph(6’)-Id,* also known as *strA/strB*; and *oqxA/oqxB*]. In addition to the “canonical (acquired) ARGs”, we also identified chromosomally located ARGs intrinsically linked to various bacterial species. These include beta-lactamases of class A (*bla*_LEN_ and *bla*_OKP_ from *Klebsiella* spp); class B (*bla*_CfiA_ from Bacteroides), class C (*bla*_MIR_ and *bla*_ACT_ from *Enterobacter*), class D (*bla*_OXA_ from *Campylobacter* and intestinal spirochaetes), and genes conferring resistance to aminoglycosides (*aac-6-Ii* from *Enterococcus faecium*), macrolides (*msrC* from *Enterococcus faecalis*), or phenicol (*catS* from streptococci). These intrinsic genes mirror the diversity of commensal bacteria in the gut of healthy individuals [48]. The diversity and distribution of ARGs varied significantly both within and between cohorts (**Figure 3, Tables S2 and S3**). Notably, Wayampi individuals showed a significantly higher average occurrence of ARGs for beta-lactam, aminoglycoside, and tetracycline (p-values < 0.05; Mann-Whitney test).

*Tetracycline resistance genes* [49,50] were the most common ARGs identified, with a significant number shared between individuals in both cohorts (41/43 allele variants of 19/24 genes, respectively, of the tetracycline resistome. Among them, 8 “core genes” were present in over 90% of the Wayampi individuals, specifically *tet(32), tet(40), tet(O), tet(Q), tet(W), tet(X*), *tetAB*(P), as mentioned above. Additionally, five genes were highly prevalent, found in 60-80% [*tet(M), tet(44)*, *tet(B)*, *tet(O/W/O)*] and 30-50%, [*tet(O/W), tet(W/32/O)*] of individuals. The remaining *tet* genes were present in less than 10% of individuals in each group. The GC content of these *tet* genes varied from 29% to 43%, suggesting different origins and varying distribution among gut microbiota. For instance, *tet(W), tet(32)* and *tet(O)* have been linked to members of Bacillota (the Lachnospiraceae family) as well as *tet(44)* (the Erysipelotrichaceae family), and *tetA(P)* (the Clostridiaceae family); *tet(Q*) and *tet(X)* are linked to various orders of the Bacteroidia class such as Bacteroidales (Bacteroideaceae, Prevotellaceae, Porphyromonadaceae), or Flavobacteriaceae (Capnocytophagaceae)[51]. A previous study indicated that these prokaryotic groups were the predominant taxa of the microbiomes of this Amerindian population [23]. All tetracycline resistance mechanisms described to date, namely ribosomal protection, efflux pumps, and enzyme modification, were associated with the *tet* genes detected in this study.

*Macrolide resistance genes* [50,52,53] were also highly diverse, with most shared between individuals from both cohorts (32/36 allele variants across 13/19 genes for Amerindians and Europeans, respectively). Similar to the *tet* genes, some macrolide resistance genes are widely distributed, suggesting a potential association with the overall composition of the human gut microbiome. All “core population genes” [*erm(B), erm(F), mef(A)*], were represented by multiple variants at high frequencies, often with several variants within the same individual sample, indicating *genetic redundancy*. However, other genes were unevenly distributed between the cohorts. For instance, *erm(G)* or *lnu*(C) were present in 90% of Europeans but in less than 10% of Wayampi. Conversely, *msr*(C), a species-specific gene of *Enterococcus faecalis,* was found in 75% of Wayampi individuals but absent in the European sample.

The pool of genes encoding *beta-lactamases* [54] includes 117 alleles across 29 gene categories. Among these, 26 are classified as genes encoding beta-lactamases of class A (Cep, Cfx, AC1, TEM, SHV, CTX-M, LEN, OXY, OKP, MAL, BlaZ), class B (CfiA), class C (ACC, ACT, CFE, CMG, CMH, DHA, CMY, MIR), and class D (OXA). Most of these genes are species-specific chromosomal genes, making them useful phylogenetic markers. In contrast to ARGs that confer resistance to tetracyclines or macrolides, only a few β-lactamase genes were highly prevalent. They included the chromosomal *bla*_ACI_ gene from *Acidaminococcus fermentans,* a species of Negativicutes, found in 50% of Wayampi individuals but only 10% of Europeans; the mobile *bla*_Cfx_ gene, which confers cephalosporinase activity in *Bacteroides* and other anaerobe species, present in 70-90% of all individuals studied, and often associated with transposable elements (Tn*4455*, Tn*7730*, Tn*7731*) [55]; and the *bla*_TEM1B_, found in 30-40% of both populations and widely distributed on Tn*3* like elements located on plasmids. While the diversity and abundance of *bla* alleles were similar between Amerindians and Europeans, their distribution differed significantly. For example, the *bla*_CepA_ gene, a cephalosporinase linked to division I of *Bacteroides fragilis*, was highly prevalent in Europeans but present in less than 10% of Amerindians. In contrast, the gut microbiomes of Amerindians were enriched with genes encoding OXA variants associated with *Brachyspira pilosicoli* (30% were producers of OXA-471, OXA-63), *Campylobacter* (such as OXA-61, -193, -460, and -461), or *Acinetobacter* (OXA-51-like), while Europeans more commonly carried OXA variants related to *Pseudomonas* (e.g. OXA-50-like). Differences in class C beta-lactamases further reflect variations in the composition of Gammaproteobacteria. Overall, most beta-lactamase genes were detected sporadically.

*Aminoglycoside resistance genes* [56] represented the second group of ARGs with high diversity, comprising 53 alleles across 31 genes. The predominant resistance mechanism was the production of aminoglycoside-modifying enzymes, either intrinsic or acquired. These enzymes include N-acetyltransferases (AAC), specifically AAC(3’’) and AAC(6’); O-phosphotransferases (APH), such as APH(2’’), APH(3’’), and APH(6’); and O-nucleotidyl or adenylyltransferases (ANT or AAD), including ANT(3’’) and ANT(6’). In addition, the acquired 16S rRNA methyltransferase (MTase) RmtF was detected sporadically. As with other ARGs, differences in gene distribution were observed between the cohorts. Genes encoding APH(3’’) and ANT(6’) enzymes were the most commonly represented in both groups, with some detected in over 90% of individuals. This included the acquired *aph*(3’’)-IIIa and *ant(*6’)-Ia genes, commonly associated with widespread MGEs in staphylococci and enterococci. Other highly prevalent genes were *aph*(3’’)-Ib (also known as *strA*), found in 60% of individuals in both groups and widespread in *Enterobacteriaceae* and *Pseudomonas*. Genes encoding enzymes of the AAC(6) family were represented by chromosomal variants specific to enterococcal species in Amerindians and by acquired plasmid-located genes in Europeans. Most of these genes were sporadic, with the exception of *aac*(6’)-Ii, a chromosomal species-specific gene of *Enterococcus faecium* found in 70% of Amerindians and 30% of Europeans. A member of the APH(6’) family, *aph(6’)-Id* (also referred to as *strB*), was present in 20-30% of both cohorts. Genes encoding enzymes of the ANT(3’’) family, which include *aad* genes traditionally associated with class I integrons, were present in less than 10% of individuals. Genes encoding APH(2) enzymes were restricted to two species-specific chromosomal genes from enterococci. Among the MTases, only RmtF was identified in Wayampi (10% of the population). Some clinically relevant genes prevalent in Europeans (30-50%) -such as *aph*(3’’)-Ia (associated with the Tn*903* and conferring Km resistance), or the bifunctional *aac(6’)-Ie-aph(2’’)* (located in Tn*4001* and conferring high level-resistance to gentamicin and kanamycin in staphylococci and enterococci), were sporadically detected in Wayampi. The high occurrence of genes linked to enterococci suggests an enrichment in this group of prokaryotes. The remaining genes, whether shared or exclusive of the two cohorts, were detected sporadically.

Genes associated with resistance to *folate inhibitors*, such as *dhfr* and *sul* genes, which are often located on MGEs, displayed significant diversity. We identified 9 *dfrA* genes, typically linked to class 1 integrons, along with *dfrG* from *Staphylococcus aureus*. Notably, *dfrA1* was the most prevalent, accounting for approximately 30% of both study cohorts, with six allelic variants in Europeans, and 16 variants in Wayampi individuals. Other *dfrA* genes were observed in about 10% of individuals from Guiana. All known *sul* genes were identified in Wayampi, with *sul2* found in over 90% of individuals from both cohorts, and *sul1* present in around 40%. The number of variants for both *sul* genes was noteworthy, particularly higher in the Wayampi group, while *sul3* was found sporadically.

*Chloramphenicol* resistance was conferred by five *cat* genes encoding chloramphenicol acetyltransferases, with prevalence ranging from 20% to 80%. The most common was the *cat* gene from *Campylobacter* (present in 80% of participants in both groups), followed by chromosomal variants from *Streptococcus*, *Clostridiales*, and Tn*9*-associated elements.

Finally, low prevalent genes were associated with resistance to *fosfomycin* (*fosA*), *glycopeptides* (*vanG*), and *quinolone* resistance (*qnrB*), detected in less than 10% of the groups.

### Diversity and distribution of metal resistance genes (the metal resistome)

MRGs represented the most abundant gene category, comprising 2411 annotated genes found in public databases. This pool included genes that help organisms to cope with the limitation of essential metals (Cu, Zn, Ni, Co, Mn, and Fe), and genes conferring resistance to highly toxic metals (Hg, As, Cd, Pb), moderately toxic metals (e.g. Co, Ni, Ag), and toxic concentrations of essential metals. In both groups of individuals, genes associated with copper resistance predominated, followed by those linked to arsenic. Genes related to zinc, iron, cobalt, and nickel genes were present though at lower and relatively similar frequencies (**Figure 2C and 2D, Table S2**).

Most MRGs identified were housekeeping genes responsible for the uptake of essential metals, and they were present in the majority of samples. The only acquired MRGs observed were related to the detoxification of copper (*pcoABCDRS* and *tcrB*), silver and copper (*silABCFPRS*), arsenic (*ars*), and mercury (*mer*). Occurrence of these genes differed between populations being higher in Europeans than Amerindians (37% vs 26% for copper, 48 vs 42% for silver, and 86 vs 76% for mercury, respectively taking as marker genes *pcoA*, *silA* and *merA*). In the following paragraphs, we will describe the diversity and the possible mechanisms of resistance of MRGs to provide insights into the epidemiological significance of MRGs in existing databases [36].

Genes related to *copper homeostasis* predominated, encompassing functions related to the import, export, and detoxification of copper. We identified four systems involved in this response (CopA/Cue, Cus, Cut, and Pco) along with a set of copper-sensing transcriptional regulators (CueR, CusR, CusS, pcoR, and pcoS), which modulate the expression of these systems. These elements represent the archetypal copper resistance response in Proteobacteria, functioning under varying conditions of oxygen availability and metal stress [57,58]. All copper-related genes, except those in the Pco system, are typically chromosomally encoded and intrinsic MRGs, and were detected in over 96% of individuals across both groups. In contrast, genes of the *pcoABCDRS* operon, exhibited variable frequency and allelic diversity. The prevalence of *pcoA, pcoB, and pcoD* was lower than *pcoE* (21-31% vs. 2% and 24-37% vs.10% in Wayampi and Europeans, respectively), suggesting heterogeneous operon arrangements (**Table S2**). A similar pattern was observed for the *silABCFPRS* operon which confers resistance to both copper and silver. The prevalence of *silA, silB, silC,* and *silP* ranged between 37–49% in Wayampi and 44–51% in Europeans, while *silE* was less frequent (26% and 17%, respectively), again indicating variation in gene content and organization. Additional copper-related influx genes identified in databases include *zupT*, and genes encoding the outer membrane protein ComC/BhsA/YcfR and its regulator ComR [59], as well as the disulfide isomerase DsbC, which increases cellular sensitivity to copper induced redox stress [60]. We also detected acquired copper resistant genes in Gram-positive bacteria included *tcrB* or *copB/mco,* typically located on plasmids in enterococci and staphylococci, respectively [61,62]. Both genes predominated among Wayampi in comparison with Europeans (65% vs 37% for *copB* and 84% vs 3% for *tcrB*).

Genes associated with zinc, nickel, cobalt, and iron homeostasis were present at similar frequencies across the two groups. Specifically, we identified systems responsible for importing free divalent cations of Zn (*zupT*, *znuABC*, and *zitB*) [63,64], Ni (*nikABCDER*) [65], Fe (*fecDC*), and Mn (*mntH/yfeP*) [64]. We also identified genes for the uptake of metal complexes such as *fecCD* (for Fe3+ and citrate) and *ybtPQ* (part of the yersiniabactin operon involved in Fe and Cu acquisition via siderophores, also referred to as metallophores). Systems involved in cobalt and nickel homeostasis included *rcnABR,* while iron-related systems included *fieF/Yyp, fetB/ybbM,* and *pmR*. Some of these transport systems show broad substrate affinity and are associated with the uptake of different metals, such as *zupT* (Zn, Fe, Co, Mn, Cu, Cd), *mntH/yefP* (Co, Cd), or *fecCD* (Ni, Co, Zn).

MRGs linked to toxic metals such as As, Hg and Cd were also present [58,66,67]. For arsenic detoxification, we detected genes such as *glpF* and the *pst* system (which facilitate the uptake of arsenite (As³) and arsenate (As), respectively), along with *arsC* (which reduces As to As³) and *arsB* (which exports As³ from the cytoplasm to the periplasm) in almost all individuals of both groups (93–100%). In contrast, genes related to mercury resistance exhibited more variable frequencies. Genes of *mer* operons (*merA, B, C, D, E, R, T*) were detected, with *merA* and *merR1* being the most prevalent (76–86%) and displaying substantial allelic diversity (17–21 *merA* variants), and *merB* and *merE* found in only 3–4% of individuals. For cadmium, genes encoding efflux pumps such as *cadD* and *czrAB* were only detected sporadically (5 vs 7 and 5% vs 0%). However, binding proteins related to the uptake of Hg and Cd (*robA*), or Hg, Cd, and Zn (*dsbABC*), were present in most Amerindians (85 and 95%, respectively). Genes linked to non-mercury divalent cations such as *zupT* and *zntA,* may also contribute to cadmium resistance. Overall, intrinsic MRGs showed high allelic diversity (**Table S2**) and redundancy at the individual level (**Table S3**).

### Diversity and distribution of biocide resistance genes

**Figures 2E and 2F** illustrate the distribution of BRGs categories and subcategories in both cohorts, which would reflect an apparent significant differences for quaternary ammonium salts and xanthenes. However, these results should be carefully interpreted because some genes are catalogued in both subcategories [36] potentially leading to misinterpretations of biocide resistance phenotypes. Many chromosomal genes such as AcrABCDEF and TolC regulate the intake of various compounds including antibiotics and were present in nearly 90% of the individuals. Species specific efflux pumps (e.g. *kdeA, kmrA, kexD* or *kpnO* from *Klebsiella* or *emrV* from *Enterobacter*) were present at variable frequencies (**Table S2**). In addition, we detected acquired genes linked to resistance and tolerance to known antiseptics, which are often co-selected alongside ARGs. These included *qacE* (33% vs 41% in Wayampi and Europeans), *qacED* -typically located on widespread class 1 integron-(20 vs 9,4%), and *qacA* (only detected in one European). Genes previously located on plasmids containing ARGs such as *sugE* and *oqxAB,* were present in in 93% and 22-29% of individuals, respectively.

### Correlation within and between ARGs, MRGs, adn BRGs pools

We further identified genes significantly overrepresented in each population across resistance categories (**Figure 4**). In Wayampi, these genes included species-specific biomarkers such as *cfxA6* and *cfxA3* from *Capnocytophaga* and other Bacteroidales, several *tet* alleles (*tetM, tetO, tetAB*), *tcrB* alleles from enterococci and *copY/tcrY* alleles from *Leuconostoc*. Genes overrepresented in Europeans were *bexA* alleles from various *Bacteroides non fragilis* species, *copY/tcrY* alleles from *Streptococcus termophillus* and macrolide genes (*lnuC, mef, ermF*) commonly associated with anaerobes. Most of these ARGs are commonly located on MGEs hosted by the aforementioned anaerobe taxa.

**Figure 4.**
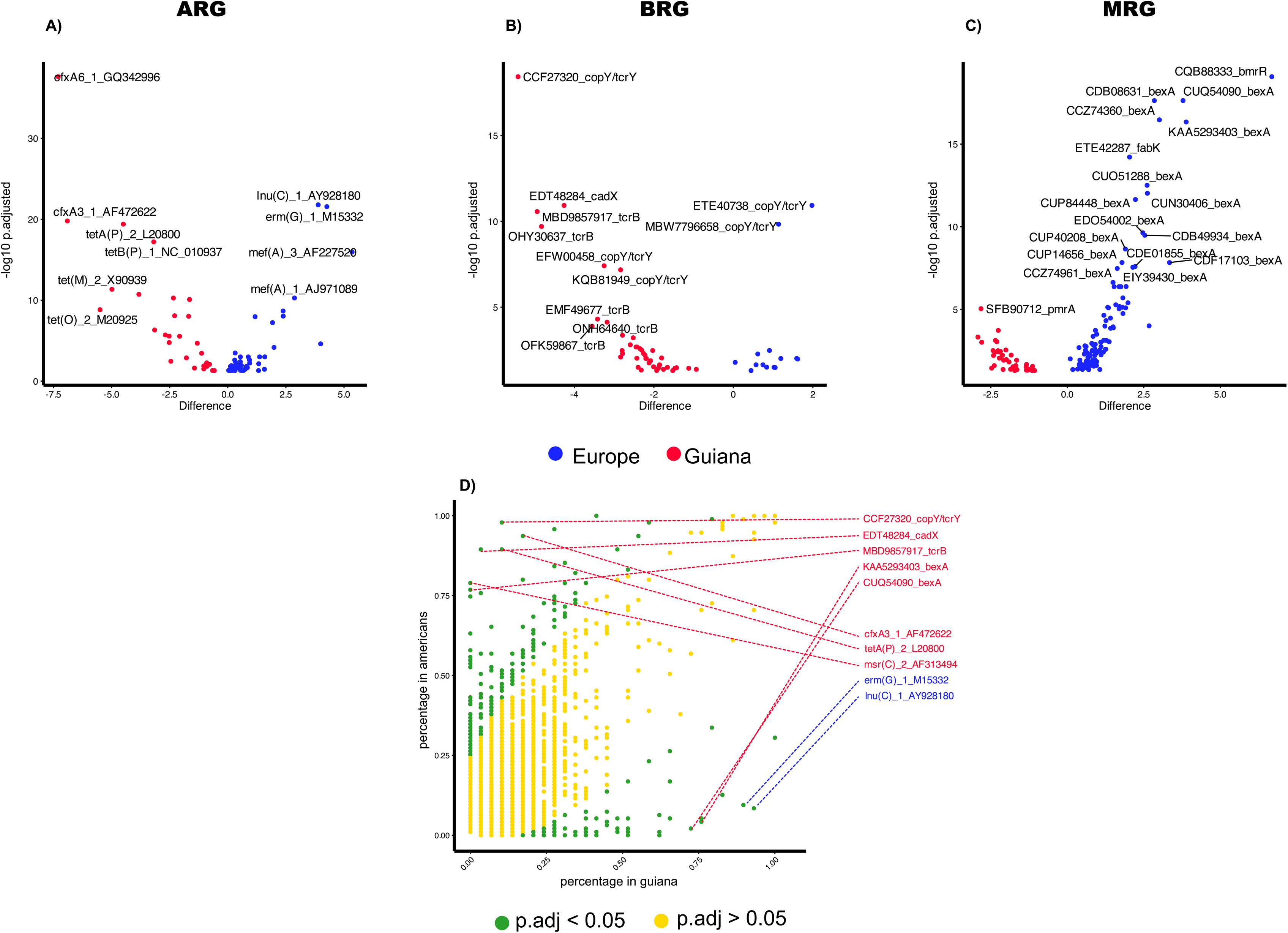
Differential abundance of resistance genes between Wayampi and European gut microbiomes. A) Volcano plots showing significantly differentially abundant genes between Wayampi (negative log2 fold-change; red) and European (positive log2 fold-change; blue) populations across ARGs, MRGs, and BRGs categories. The x-axis represents the effect size (Difference in normalized abundance), and the y-axis indicates statistical significance (-log10 of adjusted *p*-values). Only genes with a *p*-value< 0.05 are shown. (B) Prevalence scatter plot showing differential prevalence of resistance genes among Wayampi individuals and urban Americans. Each dot represents the prevalence of a resistance gene (proportion of individuals carrying the gene) in Wayampi individuals (x-axis) versus Europeans (y-axis). Genes with significantly different prevalence (p-value < 0.05) are shown in green; non-significant genes (p-value > 0.05) are shown in yellow. Red lines and labels indicate genes that are significantly more prevalent in Europeans, while blue lines and labels indicate genes that are more prevalent in the Wayampi.

Correlation analysis between genes of different categories (ARGs, MRGs, and BRGs) are shown in **Figure 5** and further detailed in the heatmaps of **Figures S3-S8** based on data from **Table S4. Figures S7 and S8** revealed distinct clustering patterns for the populations analyzed, with stronger gene correlations observed in the Europeans, particularly involving various ARGs. Notably, similar clusters observed in both groups suggest the circulation of highly conserved MGEs linked to canonical acquired genes across bacterial human populations. As examples, we identified significant association between the *pcoABCDERS* and the *silABCDEF* operons, class 1 integron gene markers (*sul1, qacED1*) and gene cassettes, plasmid-borne ARGs (*tet(C)*), species-specific efflux pumps (*smrvA/emrB* from *Enterobacter*), and species-specific beta-lactamase genes (*bla*_OKP_ from *Klebsiella, bla*_CMH_ and *bla*_MIR_ from *Enterobacter*). These associations suggest plasmids carrying these resistance genes are circulating among *Enterobacteriaceae*. The lack of correlation between the *sil* and *pco* operons supports that *sil* clusters may also circulate independently, as previously proposed.

**Figure 5.**
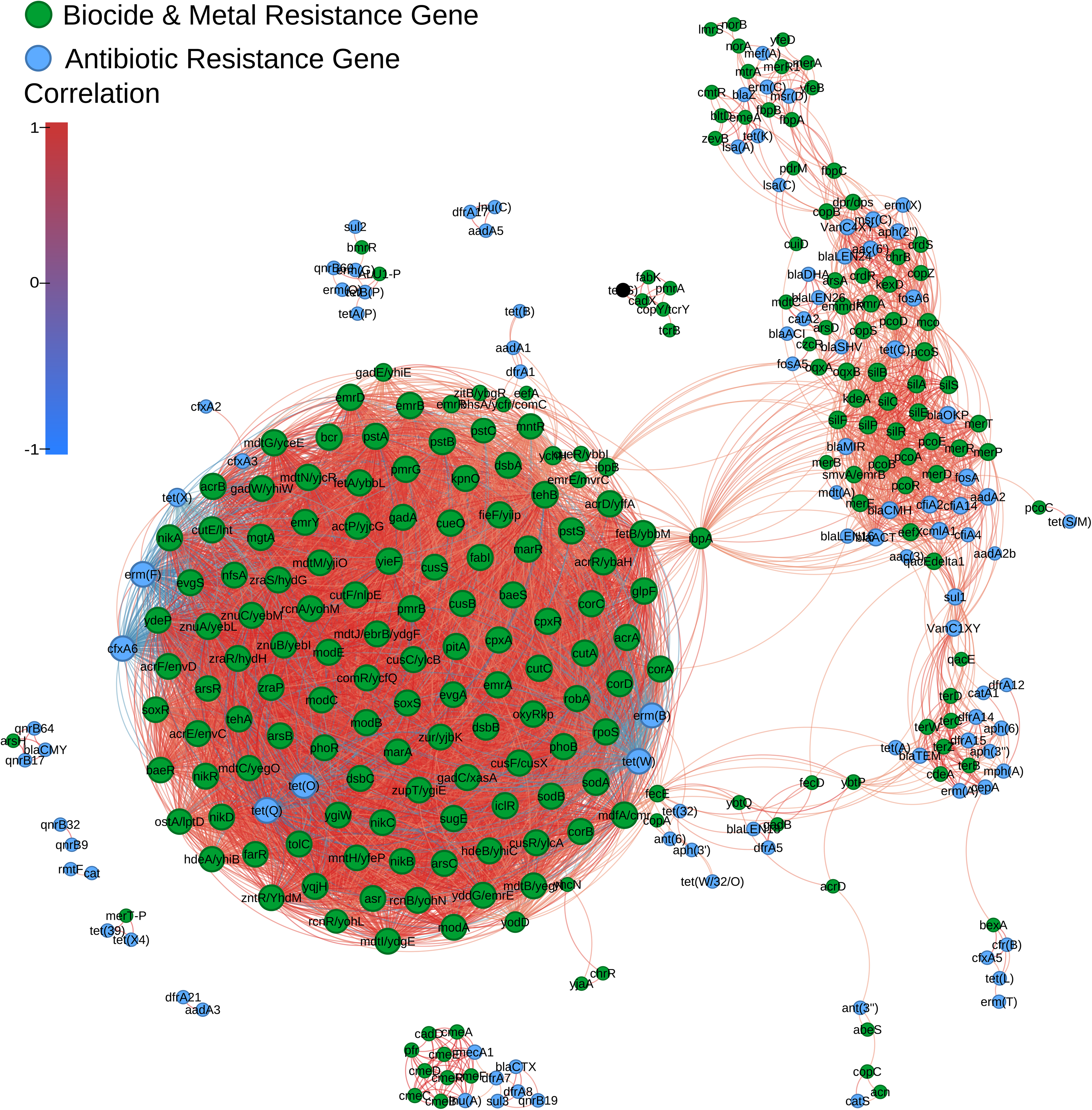
Correlation network of antibiotic, metal, and biocide resistance genes in the gut microbiome of the Wayampi population. Network representation of co-occurrence correlations among resistance genes based on Spearman’s ρ. Nodes represent individual genes, with green nodes corresponding to metal resistance genes (MRGs) and biocide resistance genes (BRGs), and blue nodes to antibiotic resistance genes (ARGs). Edges represent the correlation between nodes. Only statistically significant correlations (abs(ρ) > 0.5 and *p-value* < 0.05) are shown. Edges were colored according to the strength and direction of correlation (green = positive; red = negative). The network layout is arranged using the ForceAtlas2 algorithm.

Distinct correlation patterns were also identified among mercury resistance genes, with *merA* and *merB* clustered separately from *merC, merE, merP,* and *merT*, consistent with the presence of different *mer* operons. Nonetheless, all *mer* genes showed significant associations with both the *pco* and *sil* systems, suggesting co-location on shared MGEs. Notably, consistent associations were also observed between gene cassettes of prevalent integrons *(dfrA1-aadA1, dfrA17-aadA5, dfrA21-aadA3*) and different correlation patterns for any of these gene cassette arrays in both populations, which suggest different gene sharing patterns. In contrast, the absence of any clustering for some widespread (core) genes (*sul2, aadA2, aph3’’dfrA8*) indicates looser linkage to specific MGEs. Remarkably, negative correlations were found between core *tet* genes and housekeeping genes of *Bacteroides* with BAC genes, which are typically intrinsic to specific taxonomic groups.

### Alfa- and Beta-diversity of human cohorts according to gene categories

To compare the diversity metrics, we first assessed whether both groups followed normal distributions. The Shapiro–Wilk normality test indicated that only the Dominance Simpson Index for MRGs was normally distributed (p-value < 0.05). Based on the Student’s t-test, there were no significant differences between the groups analyzed. We then examined the diversity of ARGs, MRGs, and BRGs. Shannon index values across all gene categories ranged from 0 to 5. Median diversity values were similar for ARGs (2.8 for Amerindians *versus* 2.7 for Europeans), while higher values were observed in Amerindians for BRGs and MRGs (2.9 *versus* 2.7 and 3.4 *versus* 3.0, respectively), with statistical significance detected only for BRGs (p-value < 0.05, Mann-Whitney test) (**Figure 6A**). Using the Chao 1 richness estimator, we identified differences in the content of MRGs (110 for Amerindians *vs*. 72 for Europeans) and BRGs (71 for Amerindians *vs*. 54 for Europeans), but not in ARGs (57 for both cohorts). Additional differences in the distribution of ARGs and BRGs were further supported by the Dominance and Evenness Simpson indices (**Figure S9, Table S1)**.

**Figure 6.**
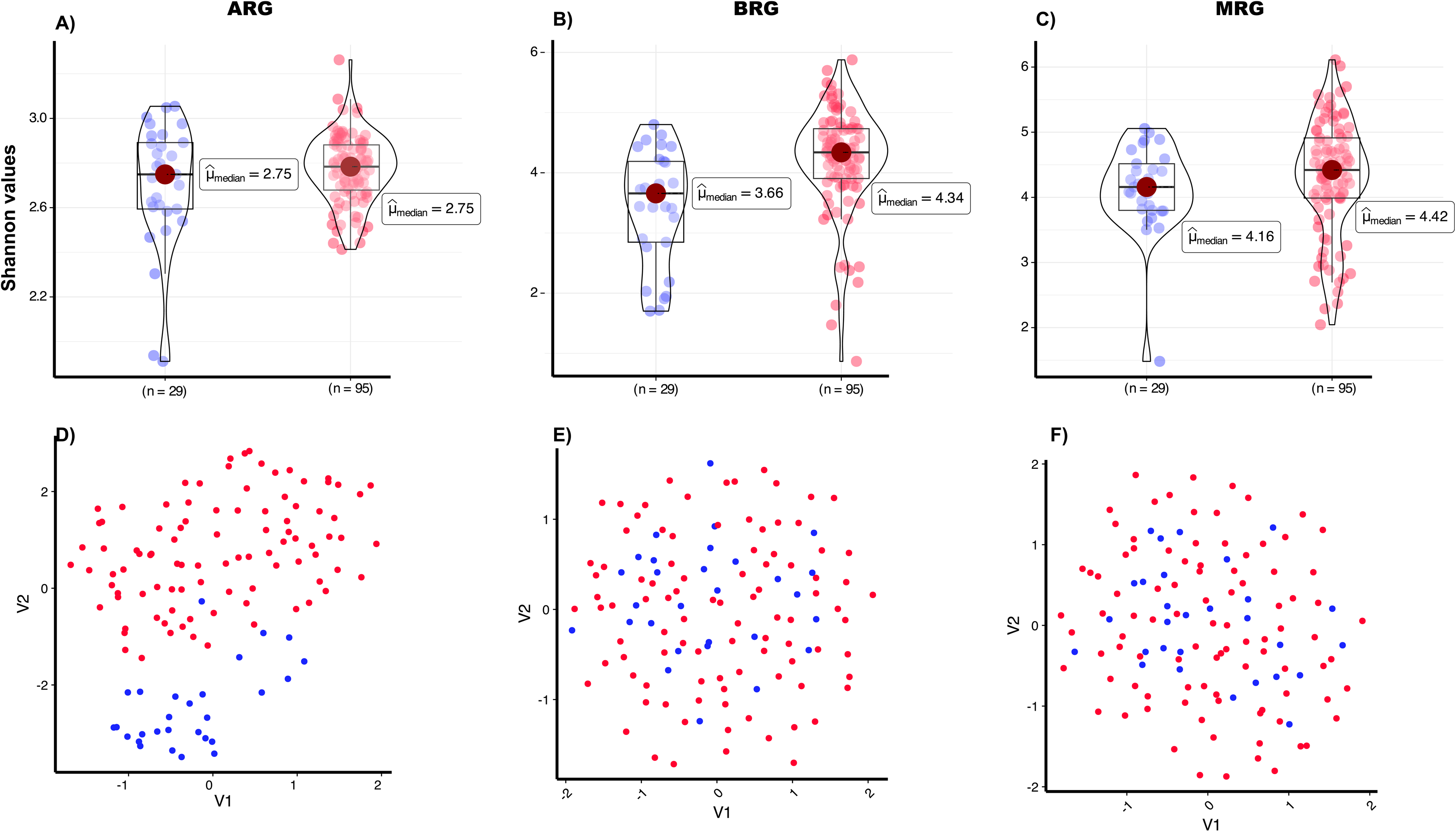
Comparison of alpha and beta diversity among Amerindian and European populations. A) Alpha diversity is assessed using the Shannon Index. The box plot combined with the violin plot illustrates the Shannon Index estimates for ARGs, MRGs, and BRGs categories. (*; p-value < 0.05) indicates statistically significant differences for BRG; (B) Uniform Manifold Approximation and Projection (UMAP) plots are used to compare resistance gene profiles across the ARGs, MRGs, and BRGs categories. The ADONIS test p-values are displayed at the top of the panel (***; p-values < 0.001). ARG=antibiotic resistance gene; MRGs=Metal resistance gene; BRG=biocide resistance gene.

To further investigate the gene composition of the different gene categories, we performed Uniform Manifold Approximation and Projection (UMAP) analysis combined with the adonis2 test (**Figure 6B**). UMAP analysis revealed distinct clustering of Amerindians and Europeans based on ARGs, but not MRGs or BRGs (ADONIS; p-values<0.001), a pattern also supported by Principal Component Analysis (PCAs) (**Figure S10**). These results indicate that while the diversity and richness of ARGs, MRGs, and BRGs pools of Amerindians and Europeans were comparable, significant differences in gene composition were detected across all categories. In addition, differences in gene evenness were observed for both ARGs and BRGs.

### Epidemiological and sociological factors associated with the Wayampiis lifestyle

Dimensionality reduction using UMAP and PCA did not reveal distinct clustering within the resistome of Wayampis. To identify variables significantly influencing the resistome composition, we performed a PERMANOVA (adonis2) analysis with 61 ecological variables and 27 variables related to antibiotic usage. For ARGs, significant associations were found with “tap water use,” “travel to the city,” “prolonged time spent outdoors,” “pregnancy,” “household size,” “type of residence,” “antibiotic use,” “fishing,” and “age.” For BACs (combined MRGs+BRGs), key factors included “tap water use,” “travel,” and “household size.” When analyzing the entire resistome, additional variables emerged, such as “vaccination within the past year,” “frequent stays outside the village,” and “use of river water.” These associations are illustrated in **Figure S11**.

## DISCUSSION

This study represents the most comprehensive characterization to date of antibiotic and non-antibiotic resistance gene pools in the human gut microbiome, using the unique model of Wayampi, an Indigenous population living in a largely pristine environment with rare antibiotic exposure but chronic environmental mercury contamination. The findings provide novel insights into the epidemiology of antimicrobial resistance that are discussed below.

Firstly, the repertoire of acquired ARGs shared between Indigenous and Westeners reveals a robust human-acquired resistome, sustained through a taxonomic gene exchange networks. The presence of a “core” set of mobile acquired ARGs (defined as genes found in all individuals regardless of lifestyle), comprising acquired genes encoding macrolides, tetracyclines and aminoglycosides, and previously observed across diverse human ecosystems [15,68], challenges the classical view that acquired ARGs may be widespread at population level though conductive gene sharing networks but not inherent to individual microbiomes [6,9]. The core mobile ARGs identified in this study have previously been detected on transposable elements linked to Bacteroidales (*tet, ermF, ermG, cfxA*), Bacteroidales and several Bacillota orders (*ermB*), or Gammaproteobacteria (*sul2, strA-strB, aphIII*) in Western human gut microbiomes [68,69], as well as in MGEs from ancient samples or minimally impacted environments [70–73]. Until recently, the origin of acquired ARGs was determined for less than 5% of unique genes in reference databases [74], mainly beta-lactamase genes that had been mobilized onto contemporary plasmids through multiple independent events [75–77]. Recent research on the tetracycline, macrolide, or aminoglycoside full resistomes [78–80] suggests early mobilization of these ARGs from diverse bacterial phyla [81,82], followed by transfer across broad phylogenetic distances. The high diversity, redundancy, and abundance of tetracycline and macrolide core genes observed in this study, long associated with dominant taxa like *Bacteroidales* and *Bacillota*, support this, indicating early mobilization, introgression (i.e., transfer to phylogenetically distinct gene pools followed by repeated backcrossing), and subsequent adaptive evolution, ultimately leading to their inheritance in certain human populations [83,84].

In addition to “core” mobile ARGs, we identified a set of shared “shell” acquired ARGs and MRGs, along with significant and conserved correlations between them. Notably, we observed consistent but patchy associations involving the *pco*, *sil*, and *mer* genes, and in some cases, ARGs linked to widespread integrons (*dfrA1-aadA1, dfrA17-aadA5, dfrA21-aadA3*), and species-specific markers such as housekeeping *bla* genes of Enterobacteriaceae (e.g. *Klebsiella* and *Enterobacter*) in both cohorts. These findings also reinforce the idea of a robust shell resistome, that while not universally present in individual gut microbiomes, is maintained at population level through a resilient mobilome [85], particularly within particular orders and genera of Gammaproteobacteria and Bacilli classes, including commensal opportunistic pathogens such as Enterobacterales, enterococci, streptococci, and staphylococci [86]. Like core genes, these shell ARGs and MRGs trace back to ancient gene clusters [87,88], increasingly detected in Enterobacterales plasmids since the mid-1950s [89–92]. Taken together, our findings support the idea of the human gut resistome as a scale-free network, with a “hub-like” core where few nodes have many connections and most nodes have very few, as previously suggested for plasmid-borne ARGs [93]. Extending this model, our results suggest the existence of distinct intermediate modules, such as species, clones, plasmids, or transposable units, that help explain the observed frequency distributions of core and shell genes across ecological gradients, which are in turn associated with taxonomically distinct bacterial and non-bacterial hosts. This structure resembles “a hourglass effect”, typical of hierarchical dependency networks [94], in which numerous inputs and outputs (such as the many antimicrobial resistance platform variants arising from HGT events) are funneled through a relatively small number of critical intermediate modules (the narrow “waist” of the hourglass) that is essential for the system’s (microbiome) function. While the maintenance, carriage and phenotypic expression of these core mobile ARGs might impose a fitness cost, many remain silent under normal conditions [95]. Others, may be regulated by invertible promoters (*invertons*)(e.g. *ermG, ermF, mefA, tetQ*) or by other regulatory mechanisms (e.g. aminoglycoside gene cassettes in integrons) in host-associated bacterial taxa, that would enable rapid adaptation of bacterial communities to disparate antimicrobial stresses [96–99].

Secondly, *gene distribution patterns differ markedly between populations* despite the existence of a common *background* resistome. Remarkable gene-level differences such as variations in genes encoding beta-lactamases from Gammaproteobacteria (OXA, AMPc) or Bacteroides (CepA, CfxA, CfiA), or macrolide resistance genes in Bacilli, suggest adaptive differences in the microbiome structure of Wayampi and Europeans. The resistome parallels the high microbial taxonomic diversity and low inter-individual variability in Wayampi, also typical of Indigenous populations, described in previous studies [22,23,100–103]. What is notable was the detection of acquired ARGs conferring resistance to first-line beta-lactam antibiotics such as *bla*_CTXM_ or *bla*_TEM-52_ which occurred following antibiotic treatment [23]. This result highlights the susceptibility of minimally impacted habitats to emerging acquired ARGs located on suitable MGEs [83] but also reveals the role of sustained antibiotic exposure (or ecological disturbance) for a broader spread and penetration of newly acquired ARGs [29], able to break the low inter-individual resistome variability. In fact, a paradoxical property of scale-free networks is a “robust-yet-fragile architecture”, highly resilient to random ecological shifts, yet susceptible to disruptions targeting central mobile genetic hubs that coordinate ARG exchange and maintenance. Our findings confirm previous observations that antibiotic resistomes are shaped by bacterial phylogeny [7], and ecology [104,105], which attributes a relevant role of the social microbiome [106].

Thirdly, this study also provides the first catalog of genes conferring resistance to antibiotics, heavy metals, and biocides in an Indigenous population. While the concept of “resistome” has largely focused almost exclusively on ARGs [2], MRGs have received comparatively little attention, despite their medical relevance [20], growing interest as alternative antimicrobials [19,20], well-documented co-selection of ARGs and MRGs [21,107,108], and environmental implications [4,109]. Most studies on MRGs have been limited to selected genes in bacterial genomes and plasmids [15,110], with few employing comprehensive metagenomic approaches [15,58]. However, the analysis of MRGs is inherently challenging due to the complexity of metal homeostasis systems, which involve genes responsible for transport, efflux, and bioconversion. These systems often display cross-affinity for multiple metals, especially first-row transition metals which share similar atomic radii and biologically relevant charges, limiting the specificity of metal-binding proteins. Although metal-binding affinities typically follow the order Zn² > Cu² > Ni² > Co² > Fe² > Mn² [111], database annotations often associated individual genes with multiple metals, increasing the risk of misinterpretation. These limitations were carefully considered in our analysis in contrast with many others that do not catalog MRGs.

The high occurrence of *mer* genes among the Wayampi aligns with prior reports of elevated mercury levels in this population, often exceeding WHO safety limits [25]. Although it is tempting to associate chronic mercury exposure through fish consumption and environmental contamination from gold mining, as suggested in our and other studies [25–27,112–115], the similar prevalence of *mer* genes in Western individuals (this study, [116]) and the robust and widespread circulation of *mer* genes in humans via mercury transposons [87], suggest an alternative or complementary explanation. Global and ancient contamination of soils, sediments, and groundwater with mercury would have enable convergent selection and further introgression of genes in different settings through the history [117]. The high conservation of certain mercury operons (transposons), *pco* and *sil* operons and their genetic contexts in human isolates supports this hypothesis. Although the metal resistome offers valuable insight into non-antibiotic antimicrobial exposures across ecosystems, its interpretation requires caution due to the functional complexity of metal resistance mechanisms, their often low specificity, current limitations in database completeness, accuracy, and taxonomic representation.

Finally, while our study addresses a critical unmet need in antimicrobial resistance research, namely, the comprehensive characterization of resistome for establishing human *background* and/or *baseline* resistomes [3,71], which is only possible through the analysis of low-impact communities, it has some limitations. These include the relatively small sample size of the European cohort and the lack of information on the genetic context of resistance genes inherent to metagenomics. Nevertheless, the primary goal of this paper was to comprehensively characterize the Wayampi resistome, and our key finding is the identification of a robust resistome that closely mirrors that of Western individuals addresses in many previous papers. The contextualizacion of our results considering the fractionated information available on both this Indigenous population (including the phylogenetic composition of their microbiome and relevant epidemiological variables [23,24,29,30]) and the current body of knowledge on antimicrobial resistance, offer insights to guide future investigations.

## CONCLUSIONS

The Wayampi resistome shows high diversity, balance, and evenness, reflecting a more less perturbed microbiome compared to industrialized populations. Yet, it reveals a shared *background* of core and shell acquired resistance genes common to human gut microbiomes. Resistance gene correlations and the swift rise of extended-spectrum beta-lactamase genes after healthcare exposure highlight a “robust-yet-fragile” ecogenetic system typical of scale free networks, resulting in vulnerability of Indigenous populations to AMR by genetically compatible genes and MGEs. The results challenge the notion that truly pristine human environments still exist [26,118] and invite a broader interpretation of *acquired antimicrobial resistance,* where the patchy yet robust distribution of acquired resistance genes reflect varying levels of conserved gene sharing developed through the co-evolution of microbial-host metacommunities.

## Supplementary Material

**Supplementary Figure S1. Rarefaction curves of resistance gene categories in the Wayampiis microbiome.** Cumulative number of antibiotic resistance genes (ARGs) identified across individuals (Left). Cumulative number of biocide and metal resistance genes (BAC = MRGs+BRGs) (Right).

**Supplementary Figure S2. Allelic gene diversity in Wayampiis.** Prevalence of resistance genes (y-axis) plotted against the number of distinct gene alleles (x-axis) across the gene categories: (A) ARGs (antibiotic resistance genes), (B) MRGs (metal resistance genes), and (C) BRGs (biocide resistance genes). Each dot represents a specific resistance gene, colored by gene family.

**Supplementary Figure S3. Heatmap of antibiotic resistance gene abundance across Wayampiis.** Rows and columns represent the abundance of ARGs detected in Wayampiis. Color intensity corresponds to relative abundance, with red indicating high abundance, yellow moderate, and blue low or no detection. Hierarchical clustering applied to both rows and columns highlights patterns of co-occurrence and differentiation among microbial communities. Data visualized using log-transformed normalized counts. Rare taxa and low-abundance noise were filtered out prior to analysis.

**Supplementary Figure S4. Heatmap of biocide resistance gene abundance across Wayampiis.** Rows and columns represent the abundance of BAC (MRGs+BRGs) detected in Wayampiis. Color intensity corresponds to relative abundance, with red indicating high abundance, yellow moderate, and blue low or no detection. Hierarchical clustering applied to both rows and columns highlights patterns of co-occurrence and differentiation among microbial communities. Data visualized using log-transformed normalized counts. Rare taxa and low-abundance noise were filtered out prior to analysis.

**Supplementary Figure S5. Heatmap of antibiotic resistance gene abundance across Europeans.** Rows and columns represent the abundance of ARGs detected in Europeans. Color intensity corresponds to relative abundance, with red indicating high abundance, yellow moderate, and blue low or no detection. Hierarchical clustering applied to both rows and columns highlights patterns of co-occurrence and differentiation among microbial communities. Data visualized using log-transformed normalized counts. Rare taxa and low-abundance noise were filtered out prior to analysis.

**Supplementary Figure S6. Heatmap of metal and biocide resistance gene abundance across Europeans.** Rows and columns represent the abundance of BAC (MRGs+BRGs) detected in Europeans. Color intensity corresponds to relative abundance, with red indicating high abundance, yellow moderate, and blue low or no detection. Hierarchical clustering applied to both rows and columns highlights patterns of co-occurrence and differentiation among microbial communities. Data visualized using log-transformed normalized counts. Rare taxa and low-abundance noise were filtered out prior to analysis.

**Supplementary Figure S7. Heatmap of resistance gene abundance across Wayampiis (antibiotic, metal, biocide).** Rows represent the abundance of ARGs and columns represent the abundance of BAC (MRGs+BRGs) detected in Europeans. Color intensity corresponds to relative abundance, with red indicating high abundance, yellow moderate, and blue low or no detection. Hierarchical clustering applied to both rows and columns highlights patterns of ARGs and BAC co-occurrence and differentiation among microbial communities. Data visualized using log-transformed normalized counts. Rare taxa and low-abundance noise were filtered out prior to analysis.

**Supplementary Figure S8. Heatmap of resistance gene abundance across Europeans (antibiotic, metal, biocide).** Rows represent the abundance of ARGs and columns represent the abundance of BAC(MRGs+BRGs) detected in Europeans. Color intensity corresponds to relative abundance, with red indicating high abundance, yellow moderate, and blue low or no detection. Hierarchical clustering applied to both rows and columns highlights patterns of ARGs and BAC (MRGs+BRGs) co-occurrence and differentiation among microbial communities. Data visualized using log-transformed normalized counts. Rare taxa and low-abundance noise were filtered out prior to analysis.

**Supplementary Figure S9. Diversity metrics of resistance genes in Amerindian and European populations**. Box plots combined with violin plots comparing diversity indices of resistance genes between European and Guianan individuals across three resistance gene categories: ARGs (antibiotic resistance genes), BACBiocide (BRGs, biocide resistance genes), and BACMetal (MRGs, metal resistance genes): **(A)** Evenness (Simpson diversity) shows gene abundance distribution; **(B)** Dominance (Simpson index) reflects the extent to which a few genes dominate the resistome; **(C)** Chao1 index estimates gene richness, accounting for rare genes. Wider plots indicate higher distribution density.

**Supplementary Figure S10. Principal Component Analysis (PCA) of resistance gene profiles in Wayampi and European individuals.** Each panel represents a different resistance gene category: ABR (antibiotic resistance genes, left), MRG (metal resistance genes, middle), and BRG (biocide resistance genes, right). Axes show the percentage of variance explained by the first two principal components.

**Supplementary Figure S11. Sociodemographic factors and predictors of resistome composition in the Wayampi population.** Heatmap showing the significance of associations between epidemiological variables (y-axis) and resistome categories (x-axis), including antibiotic resistance genes (ARGs), metal (MRGs) + biocide resistance genes (BRGs) =BAC, and antimicrobial resistance genes (ALL=ARGs+MRGs+BRGs). The color scale represents the p-values from generalized linear models, with darker shades indicating stronger statistical significance.

**Supplementary Table S1**. Statistical results associated with all analyses. The corresponding analysis is indicated in the sheet name.

**Supplementary Table S2**. Gene diversity distribution among individuals (ARGs and BAC).

**Supplementary Table S3**. Gene allelic distribution per individual (ARGs and BAC genes).

**Supplementary Table S4.** Gene correlations.

## DECLARATIONS

### Ethics approval and consent to participate

The corresponding Ethics Committees approved the enrolment of individuals from both cohorts (Comité de Protection des Personnes SudOuest et Outre Mer III, 2010-A00682-37.

### Availability of data and material

All sequences have been submitted to the EBI database under project accession number PRJEB86623.

### Authors’ contributions

M.D.F.dB.: bioinformatic and statistical analysis and manuscript writing; A.E.P.C.: statistical analysis, manuscript writing; A.A.: Wayampiís recruitment, sampling, and manuscript revision; J.L.M.: results discussion, manuscript revision; F.B.: results discussion, manuscript revision; V.F.L.: study design, data analysis, bioinformatic design, manuscript writing; T.M.C.: study design, data analysis, and manuscript writing. All the authors have read and approved the final version of this document.

## References

1. Coque TM, Cantón R, Pérez-Cobas AE, Fernández-de-Bobadilla MD, Baquero F. Antimicrobial Resistance in the Global Health Network: Known Unknowns and Challenges for Efficient Responses in the 21st Century. Microorganisms. 2023;11:1050.

2. Perry JA, Westman EL, Wright GD. The antibiotic resistome: what’s new? Curr Opin Microbiol. 2014;21:45–50.

3. Larsson DGJ, Andremont A, Bengtsson-Palme J, Brandt KK, de Roda Husman AM, Fagerstedt P, et al. Critical knowledge gaps and research needs related to the environmental dimensions of antibiotic resistance. Environ Int. 2018;117:132–8.

4. UNEP. Bracing for Superbugs: Strengthening environmental action in the One Health response to antimicrobial resistance. UNEP - UN Environ. Programme. 2023 Available from: http://www.unep.org/resources/superbugs/environmental-action.

5. Hernando-Amado S, Coque TM, Baquero F, Martínez JL. Defining and combating antibiotic resistance from One Health and Global Health perspectives. Nat Microbiol. 2019;4:1432–42.

6. Martínez JL. Natural antibiotic resistance and contamination by antibiotic resistance determinants: the two ages in the evolution of resistance to antimicrobials. Front Microbiol. 2012;3:1.

7. Forsberg KJ, Patel S, Gibson MK, Lauber CL, Knight R, Fierer N, et al. Bacterial phylogeny structures soil resistomes across habitats. Nature. 2014;509:612–6.

8. Blanco P, Hernando-Amado S, Reales-Calderon JA, Corona F, Lira F, Alcalde-Rico M, et al. Bacterial Multidrug Efflux Pumps: Much More Than Antibiotic Resistance Determinants. Microorganisms. 2016;4:14.

9. Martínez JL, Coque TM, Baquero F. What is a resistance gene? Ranking risk in resistomes. Nat Rev Microbiol. 2015;13:116–23.

10. Forslund K, Sunagawa S, Coelho LP, Bork P. Metagenomic insights into the human gut resistome and the forces that shape it. BioEssays. 2014;36:316–29.

11. Jørgensen PS, Aktipis A, Brown Z, Carrière Y, Downes S, Dunn RR, et al. Antibiotic and pesticide susceptibility and the Anthropocene operating space. Nat Sustain. 2018;1:632–41.

12. Lee K, Kim D-W, Lee D-H, Kim Y-S, Bu J-H, Cha J-H, et al. Mobile resistome of human gut and pathogen drives anthropogenic bloom of antibiotic resistance. Microbiome. 2020;8:2.

13. Zhu Y-G, Gillings M, Simonet P, Stekel D, Banwart S, Penuelas J. Microbial mass movements. Science. 2017;357:1099–100.

14. Maestre-Carballa L, Navarro-López V, Martinez-Garcia M. A Resistome Roadmap: From the Human Body to Pristine Environments. Front Microbiol. 2022;13.

15. Pal C, Bengtsson-Palme J, Kristiansson E, Larsson DGJ. The structure and diversity of human, animal and environmental resistomes. Microbiome. 2016;4:54.

16. Guitor AK, Yousuf EI, Raphenya AR, Hutton EK, Morrison KM, McArthur AG, et al. Capturing the antibiotic resistome of preterm infants reveals new benefits of probiotic supplementation. Microbiome. 2022;10:136.

17. Lanza VF, Baquero F, Martínez JL, Ramos-Ruíz R, González-Zorn B, Andremont A, et al. In-depth resistome analysis by targeted metagenomics. Microbiome. 2018;6:11.

18. Rothrock Jr. MJ, Keen PL, Cook KL, Durso LM, Franklin AM, Dungan RS. How Should We Be Determining Background and Baseline Antibiotic Resistance Levels in Agroecosystem Research? J Environ Qual. 2016;45:420–31.

19. Frei A, Verderosa AD, Elliott AG, Zuegg J, Blaskovich MAT. Metals to combat antimicrobial resistance. Nat Rev Chem. 2023;7:202–24.

20. Turner RJ. The good, the bad, and the ugly of metals as antimicrobials. BioMetals. 2024;37:545–59.

21. Baker-Austin C, Wright MS, Stepanauskas R, McArthur JV. Co-selection of antibiotic and metal resistance. Trends Microbiol. 2006;14:176–82.

22. Clemente JC, Pehrsson EC, Blaser MJ, Sandhu K, Gao Z, Wang B, et al. The microbiome of uncontacted Amerindians. Sci Adv. 2015;1:e1500183.

23. Gosalbes MJ, Vázquez-Castellanos JF, Angebault C, Woerther P-L, Ruppé E, Ferrús ML, et al. Carriage of Enterobacteria Producing Extended-Spectrum β-Lactamases and Composition of the Gut Microbiota in an Amerindian Community. Antimicrob Agents Chemother. 2016;60:507–14.

24. Grenet K, Guillemot D, Jarlier V, Moreau B, Dubourdieu S, Ruimy R, et al. Antibacterial Resistance, Wayampis Amerindians, French Guyana. Emerg Infect Dis. 2004;10:1150–3.

25. Fréry N, Maury-Brachet R, Maillot E, Deheeger M, de Mérona B, Boudou A. Gold-mining activities and mercury contamination of native amerindian communities in French Guiana: key role of fish in dietary uptake. Environ Health Perspect. 2001;109:449–56.

26. Taux K, Kraus T, Kaifie A. Mercury Exposure and Its Health Effects in Workers in the Artisanal and Small-Scale Gold Mining (ASGM) Sector—A Systematic Review. Int J Environ Res Public Health. 2022;19:2081.

27. Gerson J. Gold Mining Is Poisoning Amazon Forests with Mercury. Sci. Am. Available at https://www.scientificamerican.com/article/gold-mining-is-poisoning-amazon-forests-with-mercury/

28. Skurnik D, Ruimy R, Ready D, Ruppe E, Bernède-Bauduin C, Djossou F, et al. Is exposure to mercury a driving force for the carriage of antibiotic resistance genes? J Med Microbiol. 2010;59:804–7.

29. Grall N, Barraud O, Wieder I, Hua A, Perrier M, Babosan A, et al. Lack of dissemination of acquired resistance to β-lactams in small wild mammals around an isolated village in the Amazonian forest. Environ Microbiol Rep. 2015;7:698–708.

30. Lescat M, Clermont O, Woerther PL, Glodt J, Dion S, Skurnik D, et al. Commensal *Escherichia coli* strains in Guiana reveal a high genetic diversity with host-dependant population structure. Environ Microbiol Rep. 2013;5:49–57.

31. Filoche G, Davy D, Guignier A, Armanville F. Constructing the French state in Guiana: The challenge of the Amerindian peoples’ mobility. Crit Int. 2017;75:71–88.

32. Laperche V, Maury-Brachet R, Blanchard F, Dominique Y, Durrieu G, Massabuau JC, et al. 2007; Répartition régionale du mercure dans les sédiments et les sédiments et les poissons de six fleuves de Guyane. Rapport BRGM-PDR05GUY02.

33. Ruppé E, Ghozlane A, Tap J, Pons N, Alvarez A-S, Maziers N, et al. Prediction of the intestinal resistome by a three-dimensional structure-based method. Nat Microbiol. 2019;4:112–23.

34. Florensa AF, Kaas RS, Clausen PTLC, Aytan-Aktug D, Aarestrup FM. ResFinder - an open online resource for identification of antimicrobial resistance genes in next-generation sequencing data and prediction of phenotypes from genotypes. Microb Genomics. 2022;8:000748.

35. McArthur AG, Waglechner N, Nizam F, Yan A, Azad MA, Baylay AJ, et al. The Comprehensive Antibiotic Resistance Database. Antimicrob Agents Chemother. 2013;57:3348.

36. Pal C, Bengtsson-Palme J, Rensing C, Kristiansson E, Larsson DGJ. BacMet: antibacterial biocide and metal resistance genes database. Nucleic Acids Res. 2014;42:D737–743.

37. Langmead B, Salzberg SL. Fast gapped-read alignment with Bowtie 2. Nat Methods. 2012;9:357–9.

38. Clausen PTLC, Aarestrup FM, Lund O. Rapid and precise alignment of raw reads against redundant databases with KMA. BMC Bioinformatics. 2018;19:307.

39. Bortolaia V, Kaas RS, Ruppe E, Roberts MC, Schwarz S, Cattoir V, et al. ResFinder 4.0 for predictions of phenotypes from genotypes. J Antimicrob Chemother. 2020;75:3491–500.

40. Colwell RK, Mao CX, Chang J. Interpolating, Extrapolating, and Comparing Incidence-Based Species Accumulation Curves. Ecology. 2004;85:2717–27.

41. Koonin EV, Wolf YI. Evolution of microbes and viruses: a paradigm shift in evolutionary biology? Front Cell Infect Microbiol. 2012;2:119.

42. Raivo Kolde. pheatmap: Pretty Heatmaps. 2019. p. 1.0.12. Available from: https://CRAN.R-project.org/package=pheatmap.

43. Oksanen, J., Blanchet, F.G., Kindt, R., Legendre, P., Minchin, P.R., O’Hara, R.B., Simpson, G.L., Solymos, P., Stevens, M.H.H. and Wagner, H. R Package Version 2.2-0. Vegan: Community Ecology Package [Internet]. 2014. Available from: https://github.com/vegandevs/vegan, https://vegandevs.github.io/vegan/.

44. Hmisc [Internet]. 2024 [cited 2025 May 3]. Available from: https://hbiostat.org/r/hmisc/.

45. Jacomy M, Venturini T, Heymann S, Bastian M. ForceAtlas2, a Continuous Graph Layout Algorithm for Handy Network Visualization Designed for the Gephi Software. PLOS One. 2014;9:e98679.

46. Liu M, Li X, Xie Y, Bi D, Sun J, Li J, et al. ICEberg 2.0: an updated database of bacterial integrative and conjugative elements. Nucleic Acids Res. 2019;47:D660–5.

47. Partridge SR, Kwong SM, Firth N, Jensen SO. Mobile Genetic Elements Associated with Antimicrobial Resistance. Clin Microbiol Rev. 2018;31:e00088–17.

48. Manor O, Dai CL, Kornilov SA, Smith B, Price ND, Lovejoy JC, et al. Health and disease markers correlate with gut microbiome composition across thousands of people. Nat Commun. 2020;11:5206.

49. Thaker M, Spanogiannopoulos P, Wright GD. The tetracycline resistome. Cell Mol Life Sci. 2009;67:419–31.

50. Roberts MC. Environmental Macrolide–Lincosamide–Streptogramin and Tetracycline Resistant Bacteria. Front Microbiol 20112;2:40.

51. Juricova H, Matiasovicova J, Kubasova T, Cejkova D, Rychlik I. The distribution of antibiotic resistance genes in chicken gut microbiota commensals. Sci Rep. 2021;11:3290.

52. Leclercq R. Mechanisms of Resistance to Macrolides and Lincosamides: Nature of the Resistance Elements and Their Clinical Implications. Clin Infect Dis. 2002;34:482–92.

53. Roberts MC. Update on macrolide–lincosamide–streptogramin, ketolide, and oxazolidinone resistance genes. FEMS Microbiol Lett. 2008;282:147–59.

54. Bush K, Bradford PA. Epidemiology of β-Lactamase-Producing Pathogens. Clin Microbiol Rev. 2020;33:e00047–19.

55. Boiten KE, Kuijper EJ, Schuele L, van Prehn J, Bode LGM, Maat I, et al. Characterization of mobile genetic elements in multidrug-resistant *Bacteroides fragilis* isolates from different hospitals in the Netherlands. Anaerobe. 2023;81:102722.

56. Ramirez MS, Tolmasky ME. Aminoglycoside modifying enzymes. Drug Resist Updat Rev Comment Antimicrob Anticancer Chemother. 2010;13:151–71.

57. Hernández-Montes G, Argüello JM, Valderrama B. Evolution and diversity of periplasmic proteins involved in copper homeostasis in gamma proteobacteria. BMC Microbiol. 2012;12:249.

58. Pal C, Asiani K, Arya S, Rensing C, Stekel DJ, Larsson DGJ, et al. Metal Resistance and Its Association With Antibiotic Resistance. Adv Microb Physiol. 2017;70:261–313.

59. Mermod M, Magnani D, Solioz M, Stoyanov JV. The copper-inducible ComR (YcfQ) repressor regulates expression of ComC (YcfR), which affects copper permeability of the outer membrane of *Escherichia coli*. Biometals Int J Role Met Ions Biol Biochem Med. 2012;25:33– 43.

60. Hiniker A, Collet J-F, Bardwell JCA. Copper stress causes an in vivo requirement for the *Escherichia coli* disulfide isomerase DsbC. J Biol Chem. 2005;280:33785–91.

61. Hasman H, Aarestrup FM. tcrB, a gene conferring transferable copper resistance in *Enterococcus faecium*: occurrence, transferability, and linkage to macrolide and glycopeptide resistance. Antimicrob Agents Chemother. 2002;46:1410–6.

62. Zapotoczna M, Riboldi GP, Moustafa AM, Dickson E, Narechania A, Morrissey JA, et al. Mobile-Genetic-Element-Encoded Hypertolerance to Copper Protects *Staphylococcus aureus* from Killing by Host Phagocytes. mBio. 2018;9:e00550–18.

63. Rensing C, Moodley A, Cavaco LM, McDevitt SF. Resistance to Metals Used in Agricultural Production. Microbiol Spectr. 2018;6:10.

64. Robinson AE, Heffernan JR, Henderson JP. The iron hand of uropathogenic Escherichia coli: the role of transition metal control in virulence. Future Microbiol. 2018;13:745–56.

65. Macomber L, Hausinger RP. Nickel Toxicity, Regulation, and Resistance in Bacteria. Stress Environ Regul Gene Expr Adapt Bact. John Wiley & Sons, Ltd; 2016, p. 1131–44.

66. Barkay T, Miller SM, Summers AO. Bacterial mercury resistance from atoms to ecosystems. FEMS Microbiol Rev. 2003;27:355–84.

67. Sharma M, Sharma S, Paavan, Gupta M, Goyal S, Talukder D, et al. Mechanisms of microbial resistance against cadmium – a review. J Environ Health Sci Eng. 2024;22:13–30.

68. van Schaik W. The human gut resistome. Philos Trans R Soc Lond B Biol Sci. 2015;370:20140087.

69. Wallace MJ, Jean S, Wallace MA, Burnham C-AD, Dantas G. Comparative Genomics of *Bacteroides fragilis* Group Isolates Reveals Species-Dependent Resistance Mechanisms and Validates Clinical Tools for Resistance Prediction. mBio. 13:e03603–21.

70. Okubo T, Ae R, Noda J, Iizuka Y, Usui M, Tamura Y. Detection of the *sul2*–*strA*–*strB* gene cluster in an ice core from Dome Fuji Station, East Antarctica. J Glob Antimicrob Resist. 2019;17:72–8.

71. Scott LC, Lee N, Aw TG. Antibiotic Resistance in Minimally Human-Impacted Environments. Int J Environ Res Public Health. 2020;17:3939.

72. Kashuba E, Dmitriev AA, Kamal SM, Melefors O, Griva G, Römling U, et al. Ancient permafrost staphylococci carry antibiotic resistance genes. Microb Ecol Health Dis. 2017;28:1345574.

73. Petrova M, Kurakov A, Shcherbatova N, Mindlin S. Genetic structure and biological properties of the first ancient multiresistance plasmid pKLH80 isolated from a permafrost bacterium. Microbiology. 2014;160:2253–63.

74. Ebmeyer S, Kristiansson E, Larsson DGJ. A framework for identifying the recent origins of mobile antibiotic resistance genes. Commun Biol. 2021;4:1–10.

75. Barlow M, Hall BG. Origin and evolution of the AmpC beta-lactamases of *Citrobacter freundii*. Antimicrob Agents Chemother. 2002;46:1190–8.

76. Cantón R, González-Alba JM, Galán JC. CTX-M Enzymes: Origin and Diffusion. Front Microbiol. 2012;3:110.

77. Jacoby GA. AmpC beta-lactamases. Clin Microbiol Rev. 2009;22:161–82.

78. Berglund F, Böhm M-E, Martinsson A, Ebmeyer S, Österlund T, Johnning A, et al. Comprehensive screening of genomic and metagenomic data reveals a large diversity of tetracycline resistance genes. Microb Genomics. 2020;6:mgen000455.

79. Lund D, Kieffer N, Parras-Moltó M, Ebmeyer S, Berglund F, Johnning A, et al. Large-scale characterization of the macrolide resistome reveals high diversity and several new pathogen-associated genes. Microb Genomics. 2022;8:000770.

80. Lund D, Coertze RD, Parras-Moltó M, Berglund F, Flach C-F, Johnning A, et al. Extensive screening reveals previously undiscovered aminoglycoside resistance genes in human pathogens. Commun Biol. 2023;6:812.

81. Waglechner N, Culp EJ, Wright GD. Ancient Antibiotics, Ancient Resistance. EcoSal Plus. 2021;9:eESP-0027-2020.

82. Wright GD. Antibiotic resistance in the environment: a link to the clinic? Curr Opin Microbiol. 2010;13:589–94.

83. McInerney JO, McNally A, O’Connell MJ. Why prokaryotes have pangenomes. Nat Microbiol. 2017;2:1–5.

84. Faith JJ, Colombel J-F, Gordon JI. Identifying strains that contribute to complex diseases through the study of microbial inheritance. Proc Natl Acad Sci. 2015;112:633–40.

85. Redondo-Salvo S, Fernández-López R, Ruiz R, Vielva L, de Toro M, Rocha EPC, et al. Pathways for horizontal gene transfer in bacteria revealed by a global map of their plasmids. Nat Commun. 2020;11:3602.

86. Beiko RG, Harlow TJ, Ragan MA. Highways of gene sharing in prokaryotes. Proc Natl Acad Sci U S A. 2005;102:14332–7.

87. Mindlin S, Minakhin L, Petrova M, Kholodii G, Minakhina S, Gorlenko Z, et al. Present-day mercury resistance transposons are common in bacteria preserved in permafrost grounds since the Upper Pleistocene. Res Microbiol. 2005;156:994–1004.

88. Staehlin BM, Gibbons JG, Rokas A, O’Halloran TV, Slot JC. Evolution of a Heavy Metal Homeostasis/Resistance Island Reflects Increasing Copper Stress in Enterobacteria. Genome Biol Evol. 2016;8:811–26.

89. Liebert CA, Hall RM, Summers AO. Transposon Tn*21*, flagship of the floating genome. Microbiol Mol Biol Rev. 1999;63:507–22.

90. Nakahara H, Ishikawa T, Sarai Y, Kondo I, Kozukue H. Mercury resistance and R plasmids in *Escherichia coli* isolated from clinical lesions in Japan. Antimicrob Agents Chemother. 1977;11:999–1003.

91. Nakahara H, Ishikawa T, Sarai Y, Kondo I, Kozukue H, Silver S. Linkage of mercury, cadmium, and arsenate and drug resistance in clinical isolates of *Pseudomonas aeruginosa*. Appl Environ Microbiol. 1977;33:975–6.

92. Osborn AM, Bruce KD, Strike P, Ritchie DA. Distribution, diversity and evolution of the bacterial mercury resistance (mer) operon. FEMS Microbiol Rev. 1997;19:239–62.

93. Fondi M, Fani R. The horizontal flow of the plasmid resistome: clues from inter-generic similarity networks. Environ Microbiol. 2010;12:3228–42.

94. Sabrin KM, Dovrolis C. The Hourglass Effect in Hierarchical Dependency Networks [Internet]. arXiv; 2018. Available from: http://arxiv.org/abs/1605.05025

95. Deekshit VK, Srikumar S. ‘To be, or not to be’—The dilemma of ‘silent’ antimicrobial resistance genes in bacteria. J Appl Microbiol. 2022;133:2902–14.

96. Jiang X, Hall AB, Arthur TD, Plichta DR, Covington CT, Poyet M, et al. Invertible promoters mediate bacterial phase variation, antibiotic resistance, and host adaptation in the gut. Science. 2019;363:181.

97. Lan F, Saba J, Qian Y, Ross T, Landick R, Venturelli OS. Single-cell analysis of multiple invertible promoters reveals differential inversion rates as a strong determinant of bacterial population heterogeneity. Sci Adv. 2023;9:eadg5476.

98. Zhang Q, Xu N, Lei C, Chen B, Wang T, Ma Y, et al. Metagenomic Insight into The Global Dissemination of The Antibiotic Resistome. Adv Sci. 2023;10:2303925.

99. Hipólito A, García-Pastor L, Blanco P, Trigo da Roza F, Kieffer N, Vergara E, et al. The expression of aminoglycoside resistance genes in integron cassettes is not controlled by riboswitches. Nucleic Acids Res. 2022;50:8566–79.

100. De Filippo C, Cavalieri D, Di Paola M, Ramazzotti M, Poullet JB, Massart S, et al. Impact of diet in shaping gut microbiota revealed by a comparative study in children from Europe and rural Africa. Proc Natl Acad Sci. 2010;107:14691–6.

101. Schnorr SL, Candela M, Rampelli S, Centanni M, Consolandi C, Basaglia G, et al. Gut microbiome of the Hadza hunter-gatherers. Nat Commun. 2014;5:3654.

102. Yatsunenko T, Rey FE, Manary MJ, Trehan I, Dominguez-Bello MG, Contreras M, et al. Human gut microbiome viewed across age and geography. Nature. 2012;486:222–7.

103. Pérez-Brocal V, Andremont A, Moya A. Isolation in small populations of Wayampi Amerindians promotes endemicity and homogenisation of their faecal virome, but its distribution is not entirely random. FEMS Microbiol Ecol. 2018;94.

104. Fondi M, Karkman A, Tamminen MV, Bosi E, Virta M, Fani R, et al. “Every Gene Is Everywhere but the Environment Selects”: Global Geolocalization of Gene Sharing in Environmental Samples through Network Analysis. Genome Biol Evol. 2016;8:1388–400.

105. Gibson MK, Forsberg KJ, Dantas G. Improved annotation of antibiotic resistance determinants reveals microbial resistomes cluster by ecology. ISME J. 2015;9:207–16.

106. Sarkar A, McInroy CJA, Harty S, Raulo A, Ibata NGO, Valles-Colomer M, et al. Microbial transmission in the social microbiome and host health and disease. Cell. 2024;187:17–43.

107. Imran M, Das KR, Naik MM. Co-selection of multi-antibiotic resistance in bacterial pathogens in metal and microplastic contaminated environments: An emerging health threat. Chemosphere. 2019;215:846–57.

108. Seiler C, Berendonk TU. Heavy metal driven co-selection of antibiotic resistance in soil and water bodies impacted by agriculture and aquaculture. Front Microbiol. 2012;3.

109. Briffa J, Sinagra E, Blundell R. Heavy metal pollution in the environment and their toxicological effects on humans. Heliyon. 2020;6:e04691.

110. Hao X, Zhu J, Rensing C, Liu Y, Gao S, Chen W, et al. Recent advances in exploring the heavy metal(loid) resistant microbiome. Comput Struct Biotechnol J. 2021;19:94–109.

111. Irving H, Williams RJP. 637. The stability of transition-metal complexes. J Chem Soc Resumed. 1953;3192–210.

112. Cheng Y, Watari T, Seccatore J, Nakajima K, Nansai K, Takaoka M. A review of gold production, mercury consumption, and emission in artisanal and small-scale gold mining (ASGM). Resour Policy. 2023;81:103370.

113. efface_mining_gold_and_mercury_pollution_in_the_guiana_shield_0.pdf. Available from: https://www.ecologic.eu/sites/default/files/publication/2015/efface_mining_gold_and_mercury_pollution_in_the_guiana_shield_0.pdf.

114. Guyana: Amerindians threatened by mercury pollution from gold mining [Internet]. mediaclip. [cited 2025 Jan 23]. Available from: https://mediaclip.ina.fr/en/i24087266-guyana-amerindians-threatened-by-mercury-pollution-from-gold-mining.html.

115. Nyholt K, Jardine TD, Villamarín F, Jacobi CM, Hawes JE, Campos-Silva JV, et al. High rates of mercury biomagnification in fish from Amazonian floodplain-lake food webs. Sci Total Environ. 2022;833:155161.

116. López-López C, Ripoll A, Fernández-de-Bobadilla MD, Lanza VF, Baquero F, Coque TM. Biocide resistance among Enterobacteriaceae from fecal samples of children living in urban polluted areas. European Conference of Clinical Microbiology and Infectious Diseases (ECCMID) 2024.

117. Aleku DL, Lazareva O, Pichler T. Mercury in groundwater – Source, transport and remediation. Appl Geochem. 2024;170:106060.

118. Hwengwere K, Paramel Nair H, Hughes KA, Peck LS, Clark MS, Walker CA. Antimicrobial resistance in Antarctica: is it still a pristine environment? Microbiome. 2022;10:71.

