## Supplemental Figures for "The Antimicrobial Gut Resistome of the Wayampi reveals a shared *background* of antibiotic and metal resistance genes with industrialized populations, underscoring the “robust-yet-fragile” architecture of human gut microbiomes"

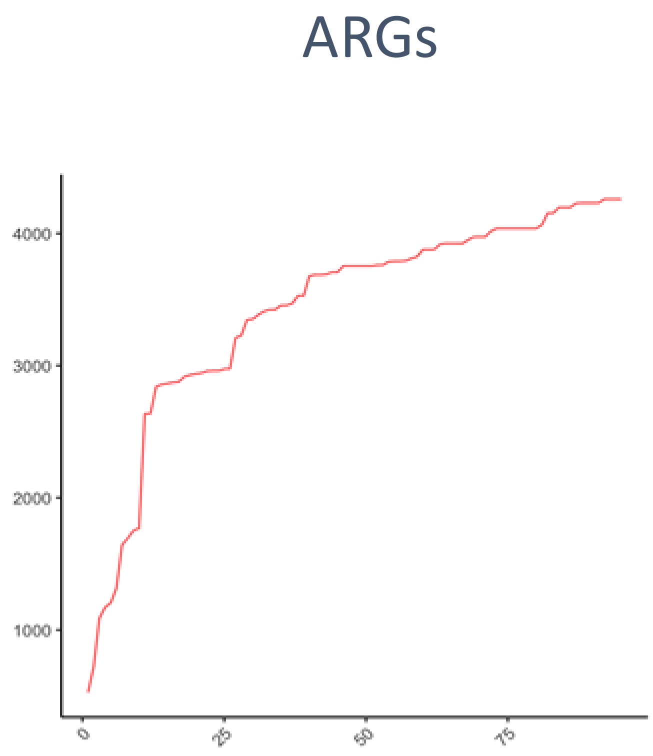
ARGs
4000
3000
2000
1000
0
25
50
75

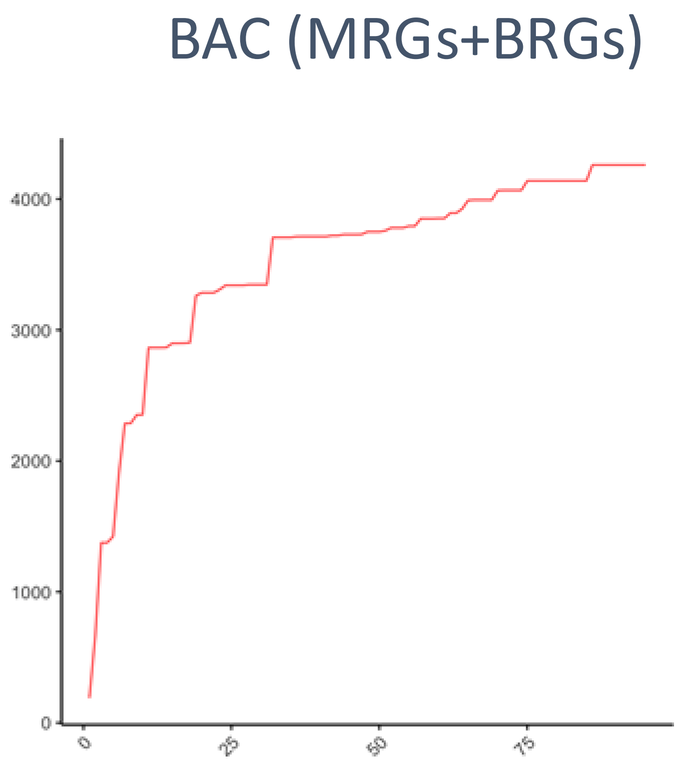
BAC (MRGs+BRGs)
4000
3000
2000
1000
0
0
25
50
75

Figure S1

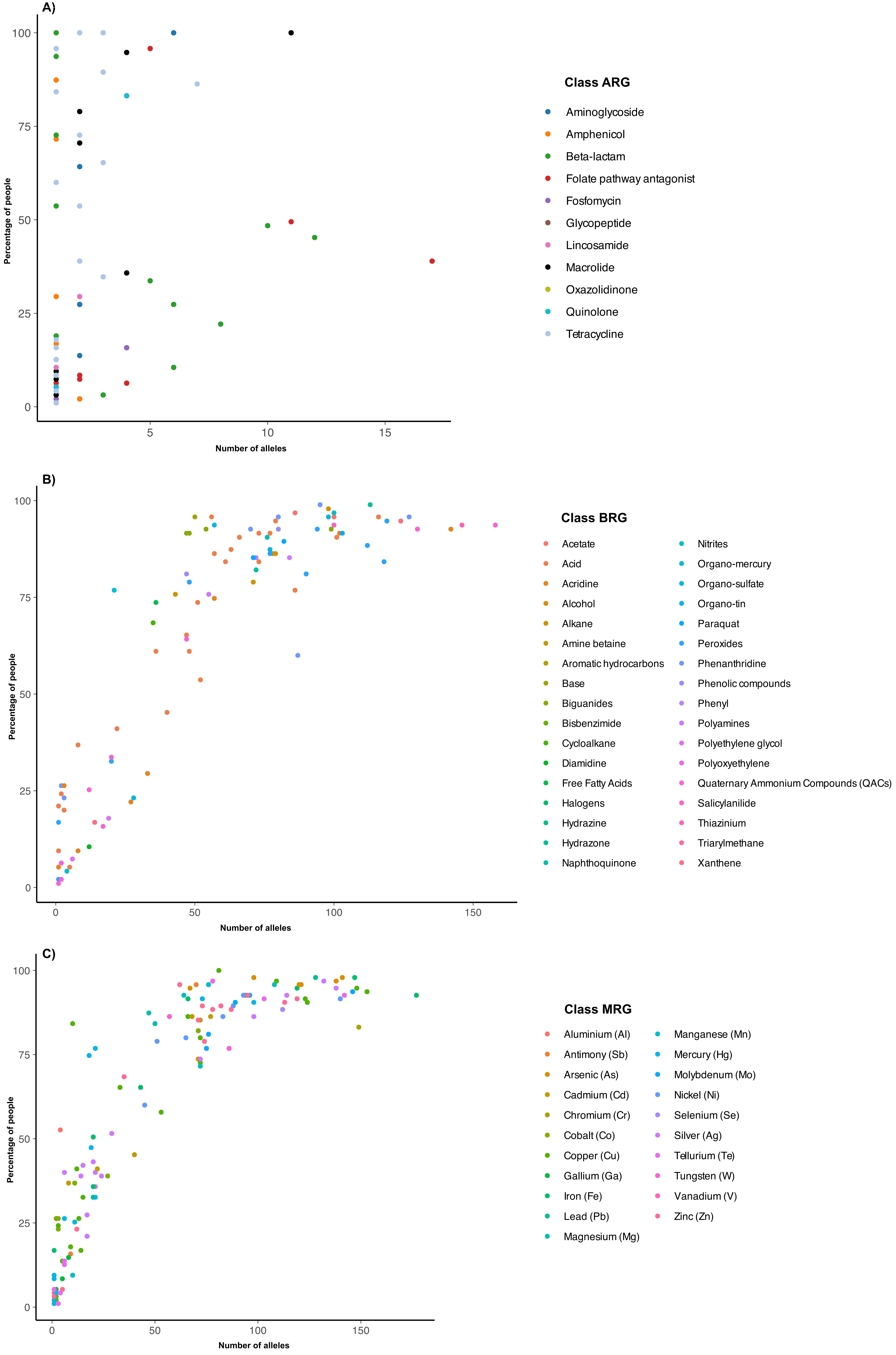
A)
Percentage of people
Number of alleles
Class ARG
Aminoglycoside
Amphenicol
Beta-lactam
Folate pathway antagonist
Fosfomycin
Glycopeptide
Lincosamide
Macrolide
Oxazolidinone
Quinolone
Tetracycline
B)
Percentage of people
Number of alleles
Class BRG
Acetate
Acid
Acridine
Alcohol
Alkane
Amine betaine
Aromatic hydrocarbons
Base
Biguanides
Bisbenzimide
Cycloalkane
Diamidine
Free Fatty Acids
Halogens
Hydrazine
Hydrazone
Naphthoquinone
Nitrites
Organo-mercury
Organo-sulfate
Organo-tin
Paraquat
Peroxides
Phenanthridine
Phenolic compounds
Phenyl
Polyamines
Polyethylene glycol
Polyoxyethylene
Quaternary Ammonium Compounds (QACs)
Salicylanilide
Thiazinium
Triarylmethane
Xanthene
C)
Percentage of people
Number of alleles
Class MRG
Aluminium (Al)
Antimony (Sb)
Arsenic (As)
Cadmium (Cd)
Chromium (Cr)
Cobalt (Co)
Copper (Cu)
Gallium (Ga)
Iron (Fe)
Lead (Pb)
Magnesium (Mg)
Manganese (Mn)
Mercury (Hg)
Molybdenum (Mo)
Nickel (Ni)
Selenium (Se)
Silver (Ag)
Tellurium (Te)
Tungsten (W)
Vanadium (V)
Zinc (Zn)

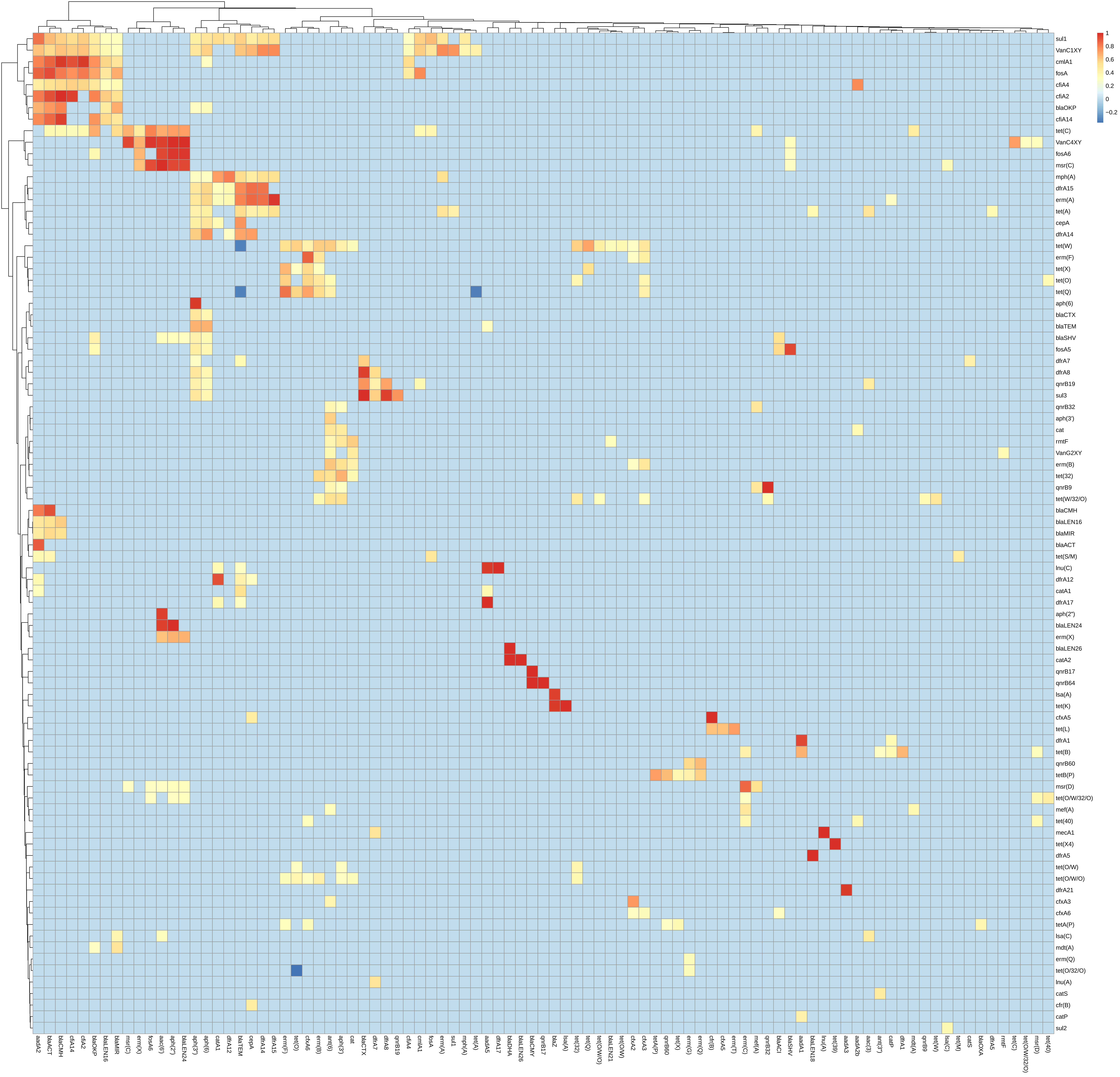
1
0.8
0.6
0.4
0.2
0
−0.2
sul1
VanC1XY
cmlA1
fosA
cfiA4
cfiA2
blaOKP
cfiA14
tet(C)
VanC4XY
fosA6
msr(C)
mph(A)
dfrA15
erm(A)
tet(A)
cepA
dfrA14
tet(W)
erm(F)
tet(X)
tet(O)
tet(Q)
aph(6)
blaCTX
blaTEM
blaSHV
fosA5
dfrA7
dfrA8
qnrB19
sul3
qnrB32
aph(3')
cat
rmtF
VanG2XY
erm(B)
tet(32)
qnrB9
tet(W/32/O)
blaCMH
blaLEN16
blaMIR
blaACT
tet(S/M)
lnu(C)
dfrA12
catA1
dfrA17
aph(2'')
blaLEN24
erm(X)
blaLEN26
catA2
qnrB17
qnrB64
lsa(A)
tet(K)
cfxA5
tet(L)
dfrA1
tet(B)
qnrB60
tetB(P)
msr(D)
tet(O/W/32/O)
mef(A)
tet(40)
mecA1
tet(X4)
dfrA5
tet(O/W)
tet(O/W/O)
dfrA21
cfxA3
cfxA6
tetA(P)
lsa(C)
mdt(A)
erm(Q)
tet(O/32/O)
lnu(A)
catS
cfr(B)
catP
sul2
aadA2
blaACT
blaCMH
cfiA14
cfiA2
blaOKP
blaLEN16
blaMIR
msr(C)
erm(X)
fosA6
aac(6')
aph(2'')
blaLEN24
aph(3'')
aph(6)
catA1
dfrA12
blaTEM
cepA
dfrA14
dfrA15
erm(F)
tet(O)
cfxA6
erm(B)
ant(6)
aph(3')
cat
blaCTX
dfrA7
dfrA8
qnrB19
cfiA4
cmlA1
fosA
erm(A)
sul1
mph(A)
tet(A)
aadA5
dfrA17
blaDHA
blaLEN26
blaCMY
qnrB17
blaZ
lsa(A)
tet(32)
tet(Q)
tet(O/W/O)
blaLEN21
tet(O/W)
cfxA2
cfxA3
tetA(P)
qnrB60
tet(X)
erm(G)
erm(Q)
cfr(B)
cfxA5
erm(T)
erm(C)
mef(A)
qnrB32
blaACI
blaSHV
aadA1
blaLEN18
lnu(A)
tet(39)
aadA3
aadA2b
aac(3)
ant(3'')
catP
dfrA1
mdt(A)
qnrB9
tet(W)
lsa(C)
tet(M)
catS
blaOXA
dfrA5
rmtF
tet(C)
tet(O/W/32/O)
msr(D)
tet(40)

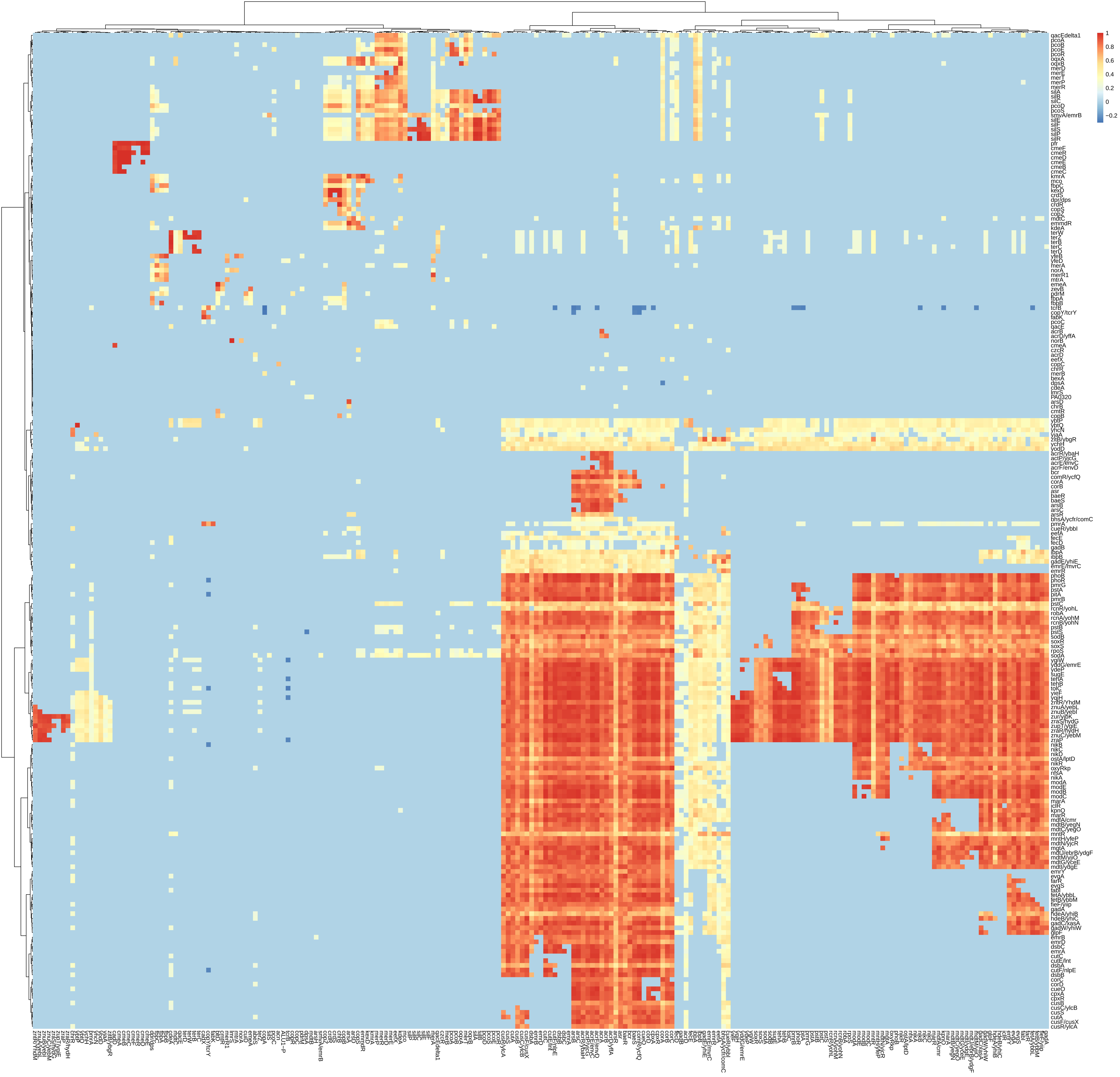
1
0.8
0.6
0.4
0.2
0
−0.2
qacEdelta1
pcoA
pcoB
pcoE
pcoR
oqxA
oqxB
merD
merE
merT
merP
merR
silA
silB
silC
pcoD
pcoS
smvA/emrB
silE
silF
silS
silP
silR
pfr
cmeF
cmeR
cmeD
cmeE
cmeB
cmeC
kmrA
mco
fbpC
kexD
crdS
dpr/dps
crdR
copS
copZ
mdtC
emmdR
kdeA
terW
terZ
terB
terC
terD
yfeB
yfeD
merA
norA
merR1
mtrA
emeA
zevB
pdrM
fbpA
fbpB
tcrB
copY/tcrY
fabK
pcoC
qacE
acrB
acrD/yffA
norB
cmeA
czcR
acrD
eefX
copC
chrR
merB
bexA
dpsA
cdeA
lmrS
PA0320
arsD
chrB
cmtR
copB
ybtP
ybtQ
yhcN
yjaA
zitB/ybgR
ychH
yodD
acrR/ybaH
actP/yjcG
acrE/envC
acrF/envD
bcr
comR/ycfQ
corA
corB
asr
baeR
baeS
arsB
arsC
arsR
bhsA/ycfr/comC
pmrA
cueR/ybbI
eefA
fecE
fecD
gadB
ibpA
ibpB
gadE/yhiE
emrE/mvrC
emrR
phoB
phoR
pmrG
pstA
pitA
pmrB
pstC
rcnR/yohL
robA
rcnA/yohM
rcnB/yohN
pstB
pstS
sodB
soxR
soxS
rpoS
sodA
ygiW
yddG/emrE
ydeP
sugE
tehA
tehB
tolC
yieF
yqjH
zntR/YhdM
znuA/yebL
znuB/yebI
zur/yjbK
zraS/hydG
zupT/ygiE
zraR/hydH
znuC/yebM
zraP
nikB
nikC
nikD
ostA/lptD
nikR
oxyRkp
nfsA
nikA
modA
modE
modB
modC
marA
iclR
kpnO
marR
mdfA/cmr
mdtB/yegN
mdtC/yegO
mntR
mntH/yfeP
mdtN/yjcR
mgtA
mdtJ/ebrB/ydgF
mdtM/yjiO
mdtG/yceE
mdtI/ydgE
emrY
evgA
farR
evgS
fabI
fetA/ybbL
fetB/ybbM
fieF/yiip
gadA
hdeA/yhiB
hdeB/yhiC
gadC/xasA
gadW/yhiW
glpF
emrB
emrD
dsbC
emrA
cutC
cutE/lnt
dsbA
cutF/nlpE
dsbB
corC
corD
cueO
cpxA
cpxR
cusB
cusC/ylcB
cusS
cutA
cusF/cusX
cusR/ylcA

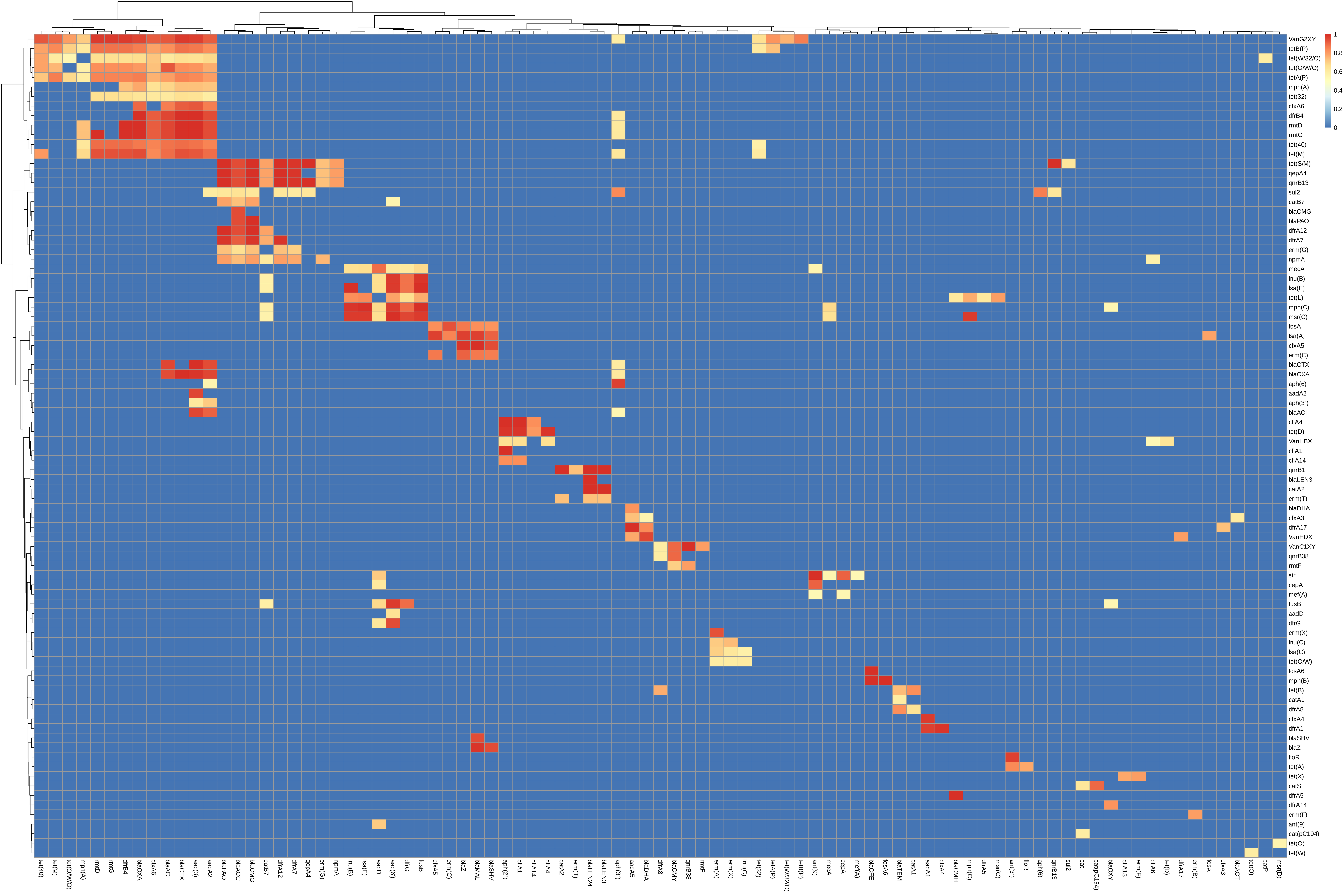
1
0.8
0.6
0.4
0.2
0
VanG2XY
tetB(P)
tet(W/32/O)
tet(O/W/O)
tetA(P)
mph(A)
tet(32)
cfxA6
dfrB4
rmtD
rmtG
tet(40)
tet(M)
tet(S/M)
qepA4
qnrB13
sul2
catB7
blaCMG
blaPAO
dfrA12
dfrA7
erm(G)
npmA
mecA
lnu(B)
lsa(E)
tet(L)
mph(C)
msr(C)
fosA
lsa(A)
cfxA5
erm(C)
blaCTX
blaOXA
aph(6)
aadA2
aph(3'')
blaACI
cfiA4
tet(D)
VanHBX
cfiA1
cfiA14
qnrB1
blaLEN3
catA2
erm(T)
blaDHA
cfxA3
dfrA17
VanHDX
VanC1XY
qnrB38
rmtF
str
cepA
mef(A)
fusB
aadD
dfrG
erm(X)
lnu(C)
lsa(C)
tet(O/W)
fosA6
mph(B)
tet(B)
catA1
dfrA8
cfxA4
dfrA1
blaSHV
blaZ
floR
tet(A)
tet(X)
catS
dfrA5
dfrA14
erm(F)
ant(9)
cat(pC194)
tet(O)
tet(W)
tet(40)
tet(M)
tet(O/W/O)
mph(A)
rmtD
rmtG
dfrB4
blaOXA
cfxA6
blaACI
blaCTX
aac(3)
aadA2
blaPAO
blaACC
blaCMG
catB7
dfrA12
dfrA7
qepA4
erm(G)
npmA
lnu(B)
lsa(E)
aadD
aac(6')
dfrG
fusB
cfxA5
erm(C)
blaZ
blaMAL
blaSHV
aph(2'')
cfiA1
cfiA14
cfiA4
catA2
erm(T)
blaLEN24
blaLEN3
aph(3'')
aadA5
blaDHA
dfrA8
blaCMY
qnrB38
rmtF
erm(A)
erm(X)
lnu(C)
tet(32)
tetA(P)
tet(W/32/O)
tetB(P)
ant(9)
mecA
cepA
mef(A)
blaCFE
fosA6
blaTEM
catA1
aadA1
cfxA4
blaCMH
mph(C)
dfrA5
msr(C)
ant(3'')
floR
aph(6)
qnrB13
sul2
cat
cat(pC194)
blaOXY
cfiA13
erm(F)
cfiA6
tet(D)
dfrA17
erm(B)
fosA
cfxA3
blaACT
tet(O)
catP
msr(D)

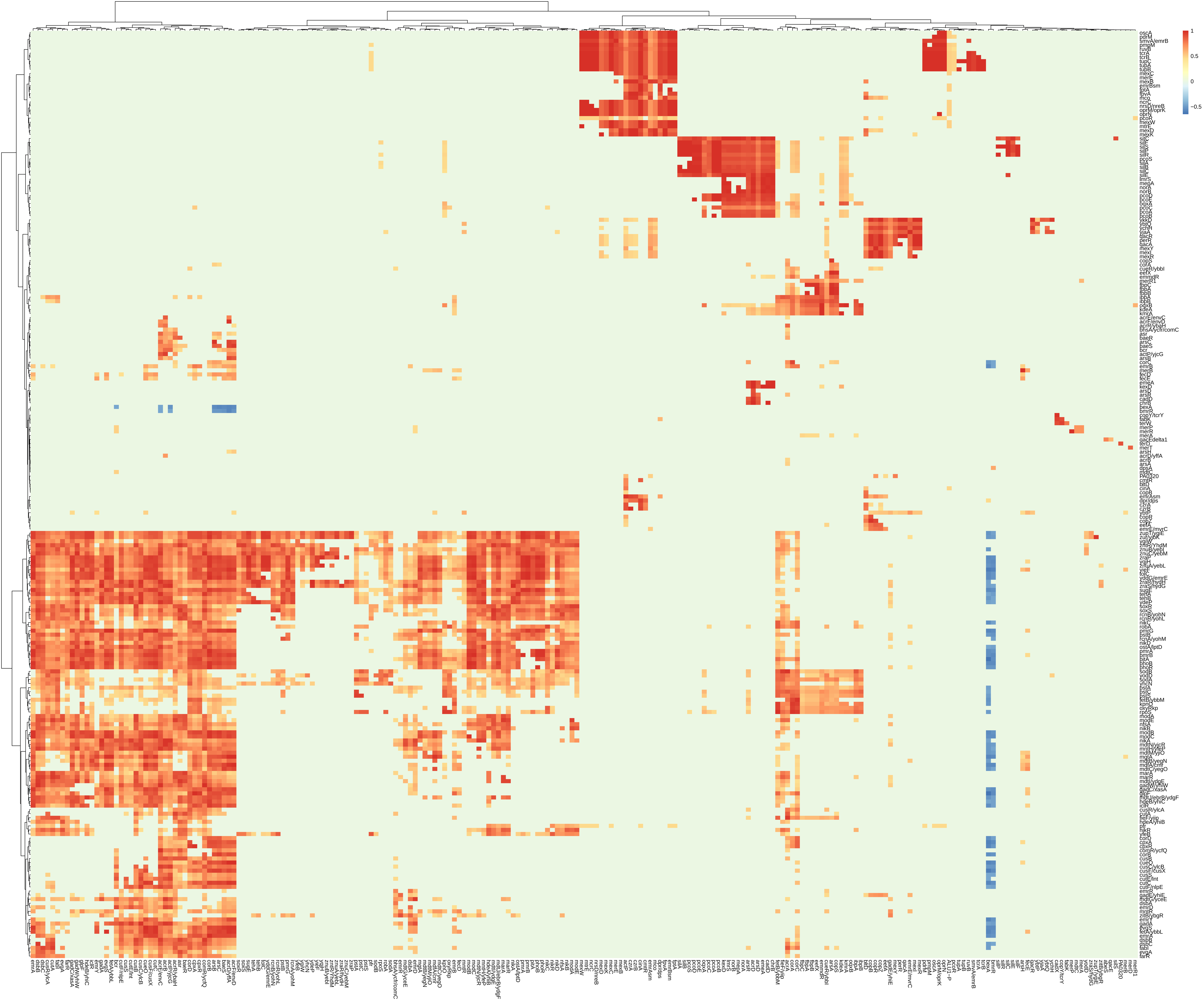
1
0.5
0
−0.5

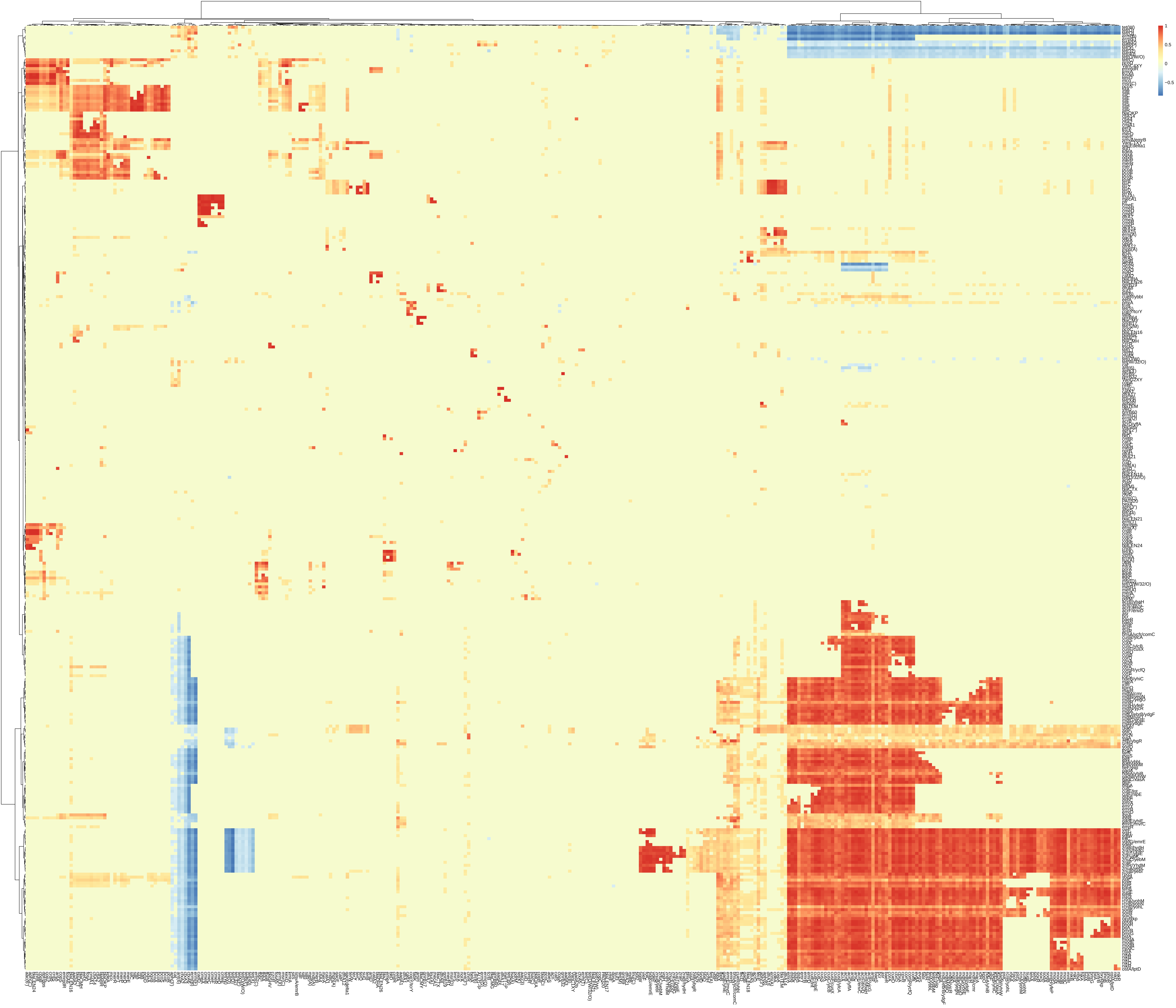
1
0.5
0
−0.5

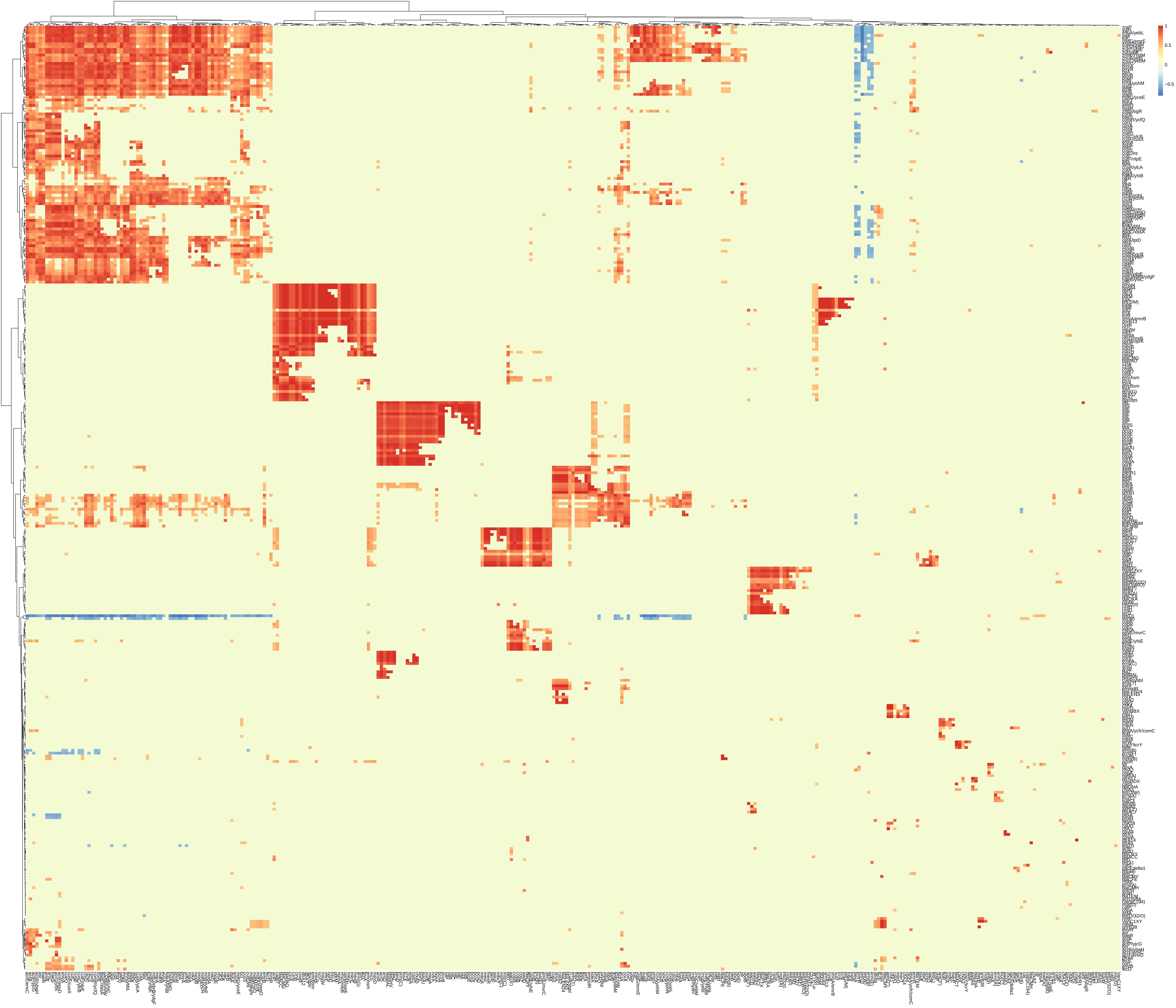
1
0.5
0
−0.5

### A) Eveness Simpson diversity

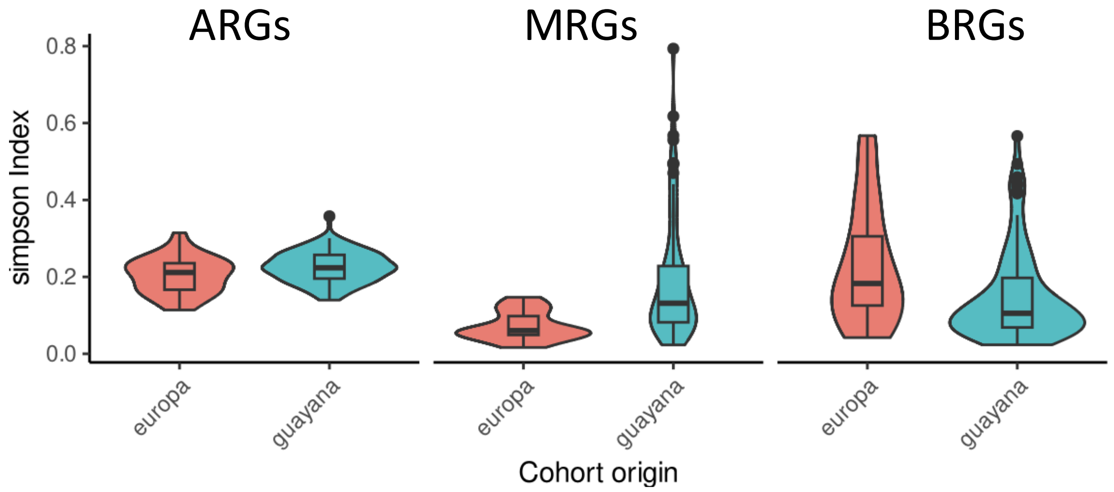


### C) Chao diversity index

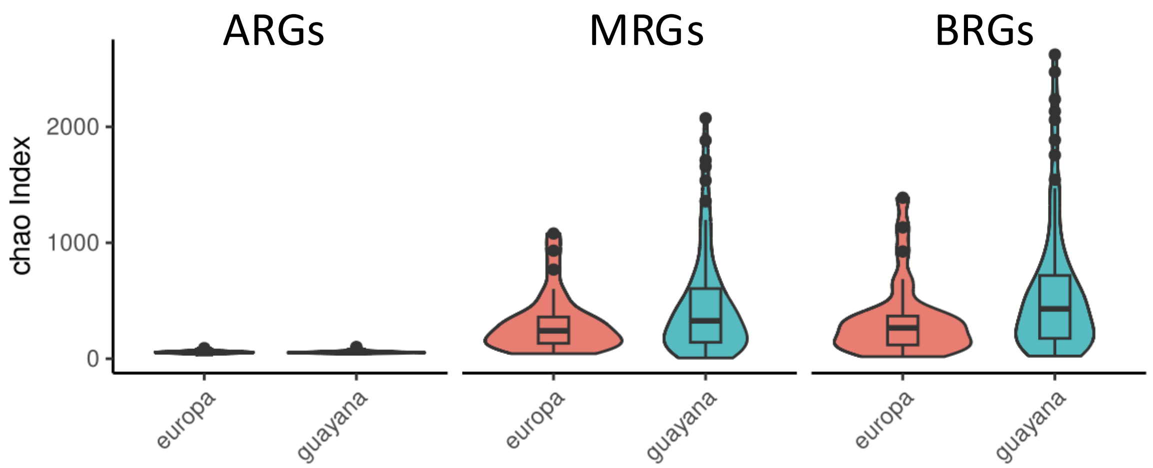


### B) Dominance Simpson diversity

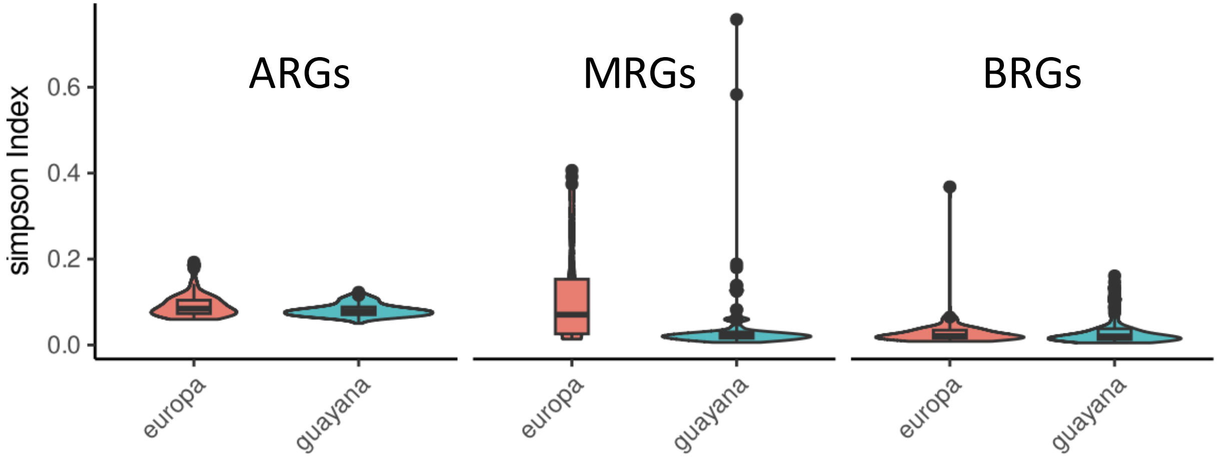

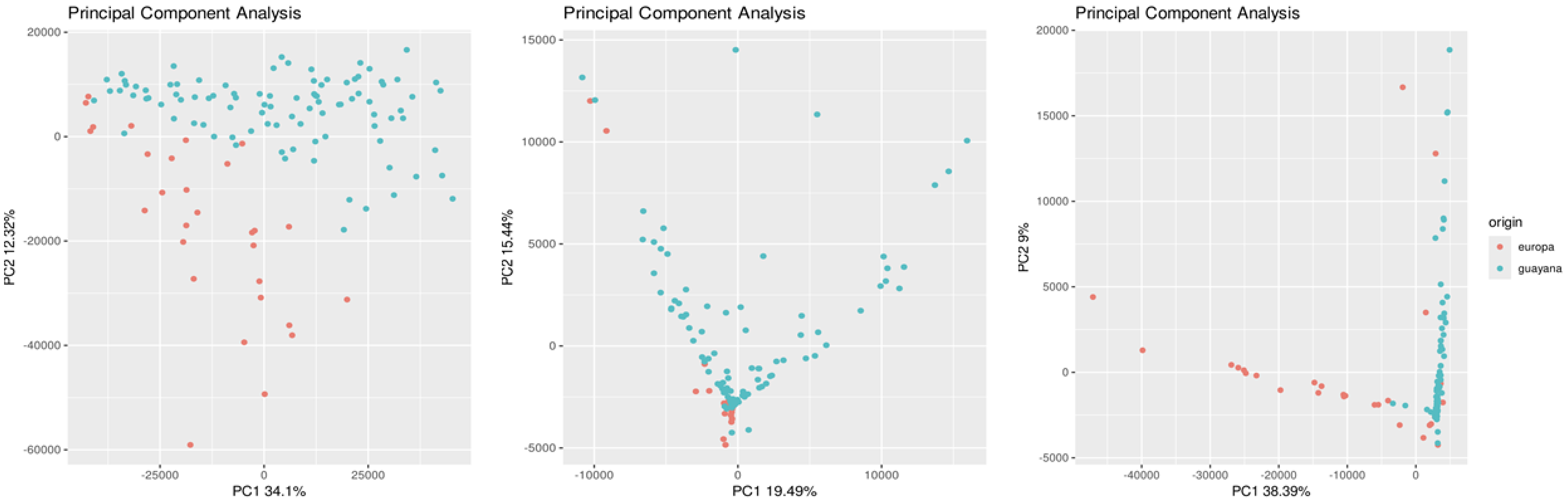
Principal Component Analysis
20000
0
-20000
-40000
-60000
-25000
0
25000
PC1 34.1%
PC2 12.32%
Principal Component Analysis
15000
10000
5000
0
-5000
-10000
0
10000
PC1 19.49%
PC2 15.44%
Principal Component Analysis
20000
15000
10000
5000
0
-5000
-40000
-30000
-20000
-10000
0
PC1 38.39%
PC2 9%
origin
europa
guayana

FigureS4

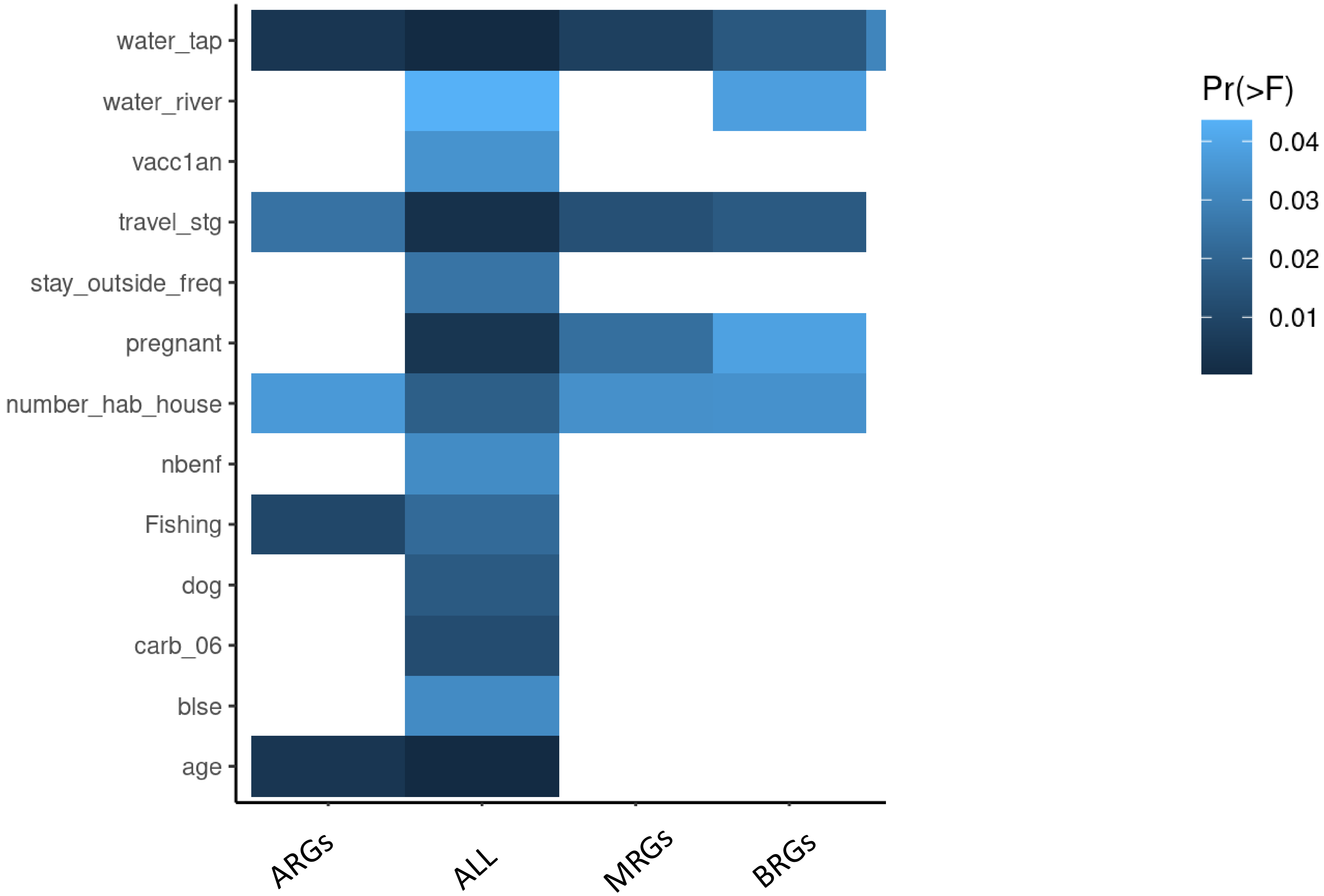
water_tap
water_river
vacc1an
travel_stg
stay_outside_freq
pregnant
number_hab_house
nbenf
Fishing
dog
carb_06
blse
age
ARGs
ALL
MRGs
BRGs
Pr(>F)
0.04
0.03
0.02
0.01

Figure S5
